# Targeting host glycolysis as a strategy for antimalarial development

**DOI:** 10.1101/2020.10.09.331728

**Authors:** Andrew J. Jezewski, Yu-Hsi Lin, Julie A. Reisz, Rachel Culp-Hill, Yasaman Barekatain, Victoria C. Yan, Angelo D’Alessandro, Florian L. Muller, Audrey R. Odom John

**Affiliations:** Department of Pediatrics, Washington University School of Medicine, St. Louis, MO; Department of Molecular Microbiology, Washington University School of Medicine, St. Louis, MO; Department of Cancer Systems Imaging, Division of Diagnostic Imaging, The University of Texas MD Anderson Cancer Center, Houston, TX; Department of Neuro-Oncology, Division of Cancer Medicine, The University of Texas MD Anderson Cancer Center, Houston, TX; Department of Biochemistry and Molecular Genetics, University of Colorado Denver – Anschutz Medical Campus, Aurora, CO; Department of Pediatrics, University of Iowa, Iowa City, IA; Department of Pediatrics, Children’s Hospital of Philadelphia, and the Perelman School of Medicine at the University of Pennsylvania, Philadelphia, PA

**Keywords:** malaria, *Plasmodium*, antimalarial, red blood cells, erythrocyte, enolase, glycolysis, oxidative stress, glutathione

## Abstract

Glycolysis controls cellular energy, redox balance, and biosynthesis. Antiglycolytic therapies are under investigation for treatment of obesity, cancer, aging, autoimmunity, and microbial diseases. Interrupting glycolysis is highly valued as a therapeutic strategy, because glycolytic disruption is generally tolerated in mammals. Unfortunately, anemia is a known dose-limiting side effect of these inhibitors and presents a major caveat to development of antiglycolytic therapies. We developed specific inhibitors of enolase – a critical enzyme in glycolysis – and validated their metabolic and cellular effects on human erythrocytes. Enolase inhibition increases erythrocyte susceptibility to oxidative damage and induces rapid and premature erythrocyte senescence, rather than direct hemolysis. We apply our model of red cell toxicity to address questions regarding erythrocyte glycolytic disruption in the context of *Plasmodium falciparum* malaria pathogenesis. Our study provides a framework for understanding red blood cell homeostasis under normal and disease states and clarifies the importance of erythrocyte reductive capacity in malaria parasite growth.

## INTRODUCTION

Infection with *Plasmodium* spp. malaria parasites contributes to over 400,000 deaths annually. Chemotherapeutic resistance remains a central roadblock to malaria control, and novel antimalarials are urgently required. One approach to circumvent resistance is through targeting host, rather than parasitic, processes essential for parasite survival. As the pathogenic stage of *Plasmodium* spp. develops within erythrocytes, the red cell niche presents a unique set of potential host targets that are needed to maintain erythrocyte metabolic homeostasis. The promise of erythrocyte-targeted antimalarial therapies is underscored by the prevalence of hereditary erythrocyte metabolic disorders in humans. These include glucose-6-phosphate dehydrogenase deficiency (G6PDd, the most common inherited enzymopathy of humans), pyruvate kinase deficiency (PKd), and sickle cell trait, all of which are under positive selection in malaria-endemic geographic regions (1–5). These prevalent genetic disorders reveal the evolutionary pressure on hosts to modulate the metabolic state of the red cell to improve survival during malaria, suggesting that acute, specific targeting of red cell glycolysis, the sole energy-generating pathway in mature erythrocytes, is a promising antimalarial approach.

Previous studies on hereditary erythrocyte enzymopathies suggest a complex interplay with malaria survival and do not provide definitive evidence as to whether acute, rather than chronic, metabolic disruption in erythrocytes indeed improves malaria clinical outcomes. For example, while reduced enzyme activity in G6PDd and PKd individuals correlates with protection from severe malaria, no specific genetic variants are associated with survival (1). In addition, severe malaria has diverse clinical presentations: while G6PDd may protect during cerebral malaria, it appears to worsen malaria-associated severe anemia, although this model is itself controversial (6, 7). Finally, acute glycolytic inhibition is distinct from the phenotype arising from hereditary enzymopathies and their complex long-term physiologic adaptations, which include increased red cell production, decreased half-life of circulating erythrocytes, and a tendency toward an overall increase of the metabolic robustness in circulating red cells (due to reticulocytosis). Similarly, caloric restriction has been proposed as a sub-acute “anti-glycolytic” state that is malaria-protective, but this intervention has pleiotropic physiologic and immunologic effects (8).

Efforts to study red cell glycolytic homeostasis using a chemical approach have been similarly complicated due to a lack of specificity. Several compounds, including sodium fluoride, arsenic, 6-aminonicatinomide, oxythiamine, and 3-bromopyruvate, are used experimentally as glycolytic inhibitors despite known off-target cellular effects (9–11). While commonly deployed as a metabolic inhibitor, 2-deoxyglucose (2-DG) is likewise complex. First, 2-DG does not completely recapitulate low glucose conditions or glycolytic inhibition, because, like glucose itself, 2-DG is processed in erythrocytes (through hexokinase and glucose 6-phosphate dehydrogenase) toward NADPH production (12–15). Second, use of 2-DG in animal models impacts all cell types, including immunocytes. Metabolic reprogramming is essential to the function of immune cells, including macrophages and natural killer cells, which themselves are well-recognized mediators of malaria-induced anemia (16–21).

Enolase (phosphopyruvate hydratase; E.C. 4.2.1.11) catalyzes the eighth step in glycolysis. Humans express three isozymes of enolase (α, β, γ) that comprise multiple dimer configurations: ENO1 (αα), found in all cell types; ENO2 (γγ), primarily in neurons; and ENO3 (ββ), a muscle-specific isoform (22). Like neurons, erythrocytes particularly rely on ENO2 for glycolytic function (23). While red bloods cells predominantly express the α isozyme they rely on the γ isozyme to a larger extent than any other non-nervous tissue (24). Thus, enolase presents a unique opportunity to achieve cellular specificity of glycolytic inhibition. In recent years, enolase inhibitors have been developed with compelling subtype specificity and enhanced potency via prodrug strategies (25–27). This newly developed class of phosphonate enolase inhibitors (some of which are ENO2-specific) and their cognate pro-drugs represent novel reagents to evaluate the role of specific and acute glycolytic disruption in erythrocytes during malaria infection. Neurons, which express both ENO1 and ENO2, withstand ENO2 specific inhibition, while erythrocytes do not (27). Here we query and define the signature of disrupted red cell glycolytic homeostasis, validate tools to investigate antiglycolytic approaches to human disease states, and use these tools to establish the impact of acute and specific glycolytic inhibition of erythrocytes on *Plasmodium* spp. infection.

## RESULTS

### Metabolic profile of acute glycolytic inhibition in human erythrocytes

Enolase (E.C. 4.2.1.11) catalyzes the penultimate glycolytic step, the conversion of 2-phosphoglycerate (2-PG) to phosphoenolpyruvate (PEP). The phosphonate natural product SF-2312 inhibits human enolase with high micromolar potency but is charged and relatively excluded from cells (25, 26). The more permeable lipophilic ester prodrugs, POM-SF and POM-HEX, are converted intracellularly to the direct enolase inhibitors SF-2312 and HEX, respectively, and have substantially increased cellular potencies over their cognate phosphonates (27). To validate POM-SF and POM-HEX as specific inhibitors of erythrocyte enolase, we performed untargeted metabolomics. Principal component analysis (PCA) and orthogonal projections of latent structures discriminant analysis (OPLS-DA) establish that POM-SF and POM-HEX-treated human erythrocytes rapidly develop distinct metabolic profiles from untreated controls. This divergence increases during prolonged exposure (Supplemental Figure 1; Supplemental Table 2-3).

Importantly, the observed metabolic profiles (Supplemental Tables 1,4) are consistent with acute and specific inhibition of glycolysis at the level of enolase, such that metabolites immediately upstream of enolase accumulate and metabolites immediately downstream are depleted (Figure 1A). Hierarchical clustering yields two prominent clusters: in the first, glycolytic inhibition slows or prevents metabolite accumulation, and in the second, glycolytic inhibition accelerates metabolite depletion (Figure 1B).

**Figure 1.**
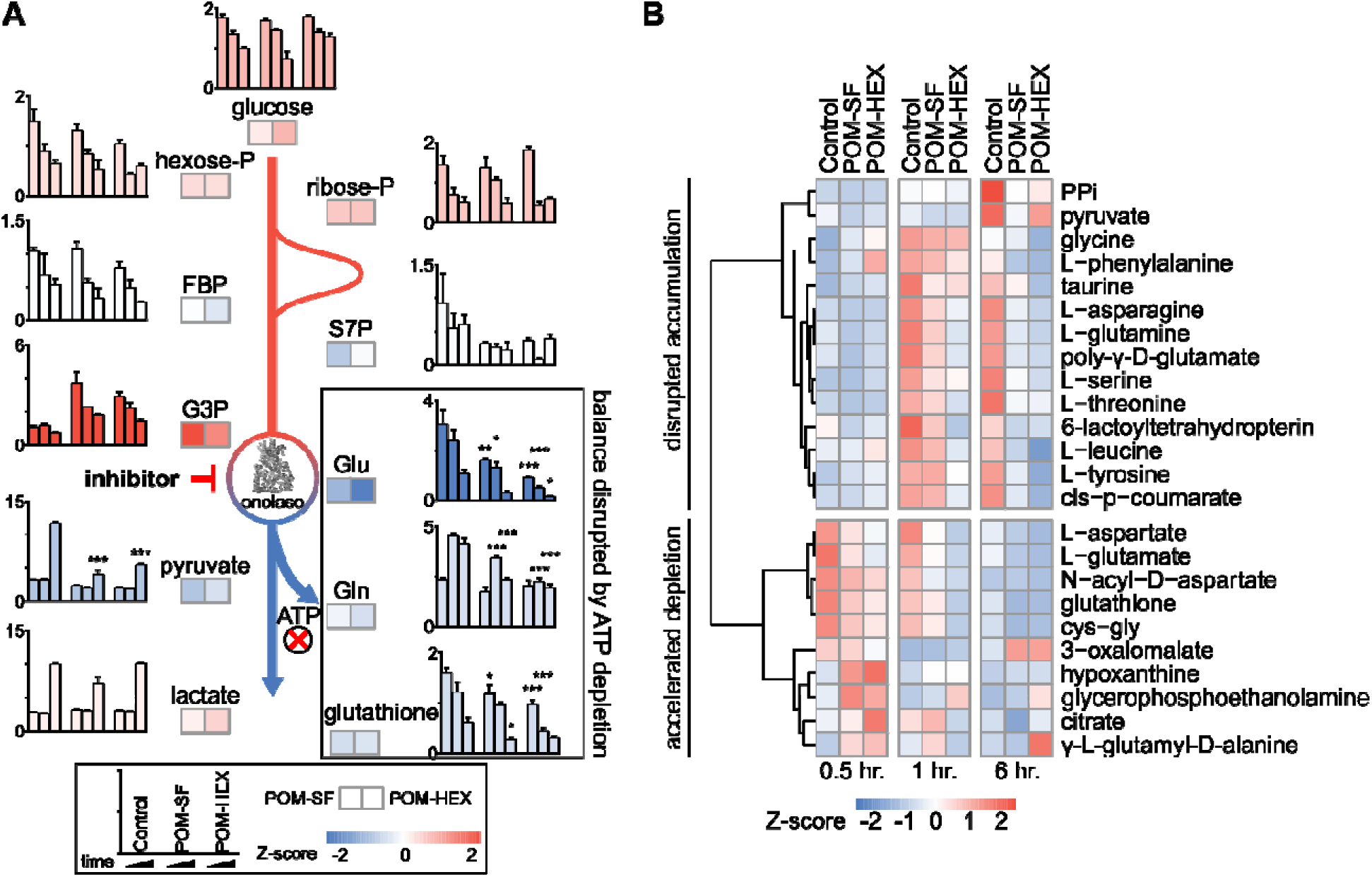
Untargeted RBC metabolomics supports on-target inhibition of enolase and establishes a signature of enolase inhibition. Enolase inhibitor treated and untreated freshly harvested RBCs were collected for metabolite detection. (A) Metabolic changes proximal and distal to enolase indicate a characteristic pattern of inhibition. Key metabolites are plotted at 0.5, 1, and 6-hr. time points normalized to 0hr. controls, for untreated, POM-SF, and POM-HEX RBCs. Z-scores of key metabolites for each drug treatment were calculated from fold changes over control samples at t = 6 hr. FBP = fructose-1,6-bisphosphate, S7P = sedoheptulose-7-phosphate, G3P = glyceraldehyde-3-phosphate. (B) Hierarchical clustering and time course for Z-scores of metabolites significantly different with respect to treatment and/or the interaction of time and treatment as calculated from the mean peak intensities from three biological replicates normalized to initial (t = 0 hr.) samples. PPi = inorganic pyrophosphate, cys-gly = L-cysteine-L-glycine. All samples were collected in experimental triplicate (n=3).

### Acute glycolytic inhibition disrupts erythrocytic glutathione homeostasis

Metabolite-pathway analysis reveals that acute inhibition of glycolysis leads to significant alterations in metabolites associated with glutathione metabolism (FDR adj. p = 4.21E-03), alanine, aspartate, glutamate metabolism (FDR adj. p = 3.97E-05), and glycine, serine, threonine metabolism (FDR adj. p = 8.34E-04) (Figure 2A; Supplemental Table 5) (28). In addition, metabolite-protein nodal analysis reveals a high degree of interconnectedness of enolase inhibitor-responsive metabolites and protein interactions. Functional analysis supports an enrichment in glutathione metabolism (p = 4.84E-52), alanine, aspartate, glutamate metabolism (p = 2.11E-34), and pyruvate metabolism (p = 7.61E-20) (Figure 2B; Supplemental Table 6). Altogether, these metabolite characterizations indicate that acute glycolytic inhibition leads to a collapse in cellular reductive capacity. As glutathione synthesis is an ATP-dependent phenomenon, inhibition of glycolysis thus decreases total red cell antioxidant capacity. In corroboration, we also find a reduction in the total glutathione pool (reduced [GSH] + oxidized [GSSG]), driven by a depletion of GSH upon enolase inhibitor treatment (Figure 1A, 2C). It is likely that the reduced availability of glutathione component amino acids further compounds this effect. As GSH is the most abundant low molecular weight cellular antioxidant, we expected that acute glycolytic disruption in erythrocytes would cause a profound susceptibility to oxidative insults.

**Figure 2.**
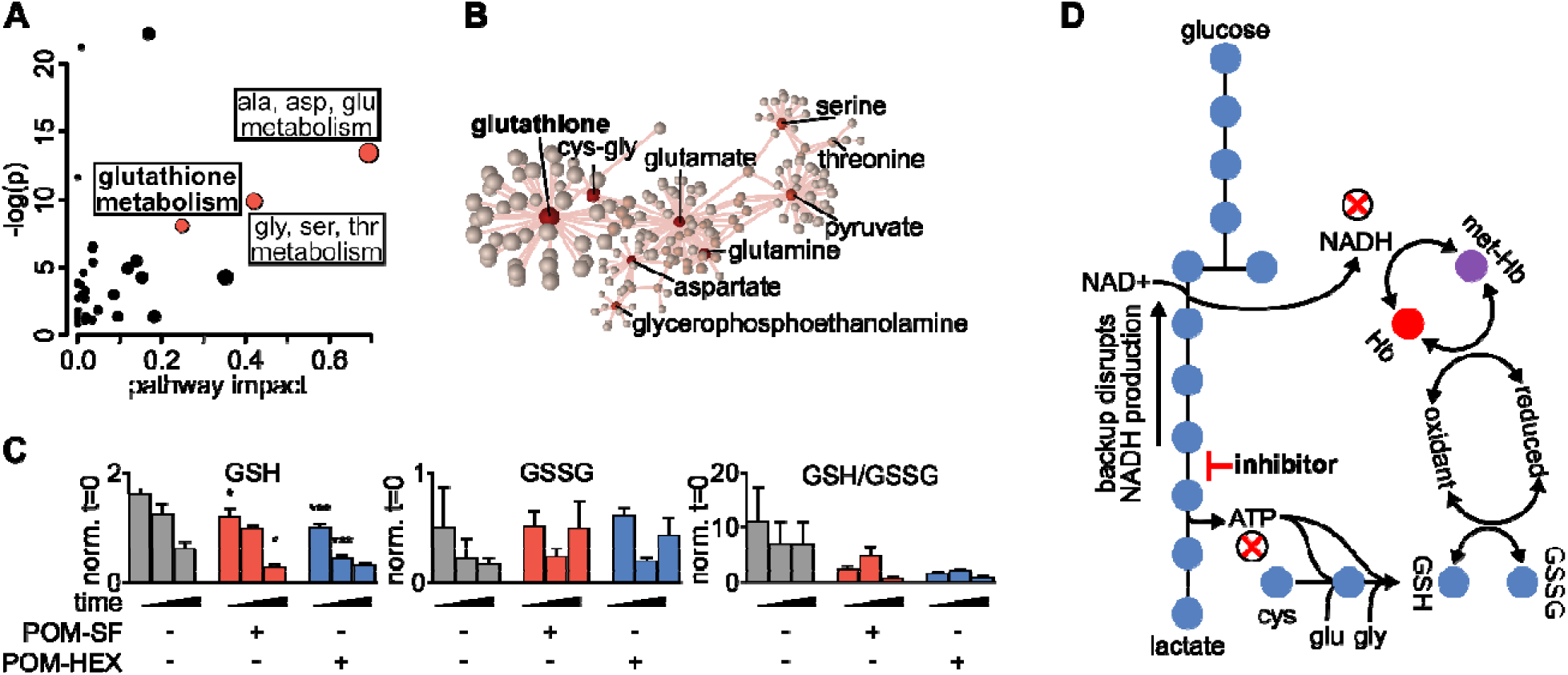
Altered glutathione metabolism in erythrocytes following inhibition of glycolysis. (A) Metabolites significantly different with respect to treatment and/or the interaction of time and treatment show an enrichment in amino acid metabolism and glutathione metabolism as highlighted in red. Pathway impact is determined by the taking the total centrality score of significantly different metabolites compared to the total centrality score of all metabolites within the pathway as described in methods. A full list of metabolic pathways associated with significantly affected metabolites and the likelihood those metabolite changes impact their associated pathway is displayed in Supplemental Table 5. (B) Nodal analysis performed as described in methods of the same metabolites as in (A) depicts a high degree of interrelatedness through their predicted metabolite-protein interactions based on the KEGG database, and this interrelatedness is enriched for glutathione production. A full list of metabolite-protein interactions is displayed in Supplemental Table 6. (C) The depletion in total glutathione is driven via a depletion in reduced glutathione during enolase inhibitor treatment. (D) Model of glycolytic inhibition disrupting glutathione metabolism. (A-C) Statistics were performed such that metabolites found to be significant following two-way repeated measures ANOVA omnibus tests adjusted for multiple comparisons using a false discovery rate of 0.05 were followed up with post-hoc t-test analyses, treatment vs. control at each time point and adjusted for multiple comparisons, Dunnett method, * = p ≤ 0.05, ** = p ≤ 0.01, *** = p ≤ 0.001. All samples were collected in experimental triplicate (n=3). cys-gly = cysteine-glycine, GSH = reduced glutathione, GSSG = oxidized glutathione, Hb = hemoglobin, met-Hb = methemoglobin.

### Acute glycolytic inhibition causes erythrocyte senescence

To evaluate oxidative susceptibility during acute glycolytic inhibition in erythrocytes, we quantified methemoglobin, in which the heme iron of hemoglobin is oxidized to the ferric (Fe^3+^), rather than ferrous (Fe^2+^) form. While methemoglobin forms spontaneously under normoxic conditions, it is typically reduced by cellular protective mechanisms, some of which are dependent on reduced glutathione (29, 30). As methemoglobin reductase is fueled by NADH (and, to a lesser extent, NADPH), we hypothesized that inhibition of late glycolysis would result in a metabolic bottleneck causing decreased NADH availability and thus impaired capacity to reduce ferric iron (Figure 3A). In keeping with this hypothesis, 1,2,3-^13^C_3_-glucose tracing highlights significant decreases in ^13^C_3_-pyruvate labeling and the ratio of ^13^C_3_-pyruvate/^13^C_3_-lactate, while flux through glycolysis versus the pentose phosphate pathway is unaltered – as gleaned by the ratios of isotopologues M+2/M+3 of lactate (Figure 3B). We observed a consistent dose-dependent increase in methemoglobin formation upon inhibitor treatment (Figure 4A), confirming a functional loss of reductive capacity.

**Figure 3.**
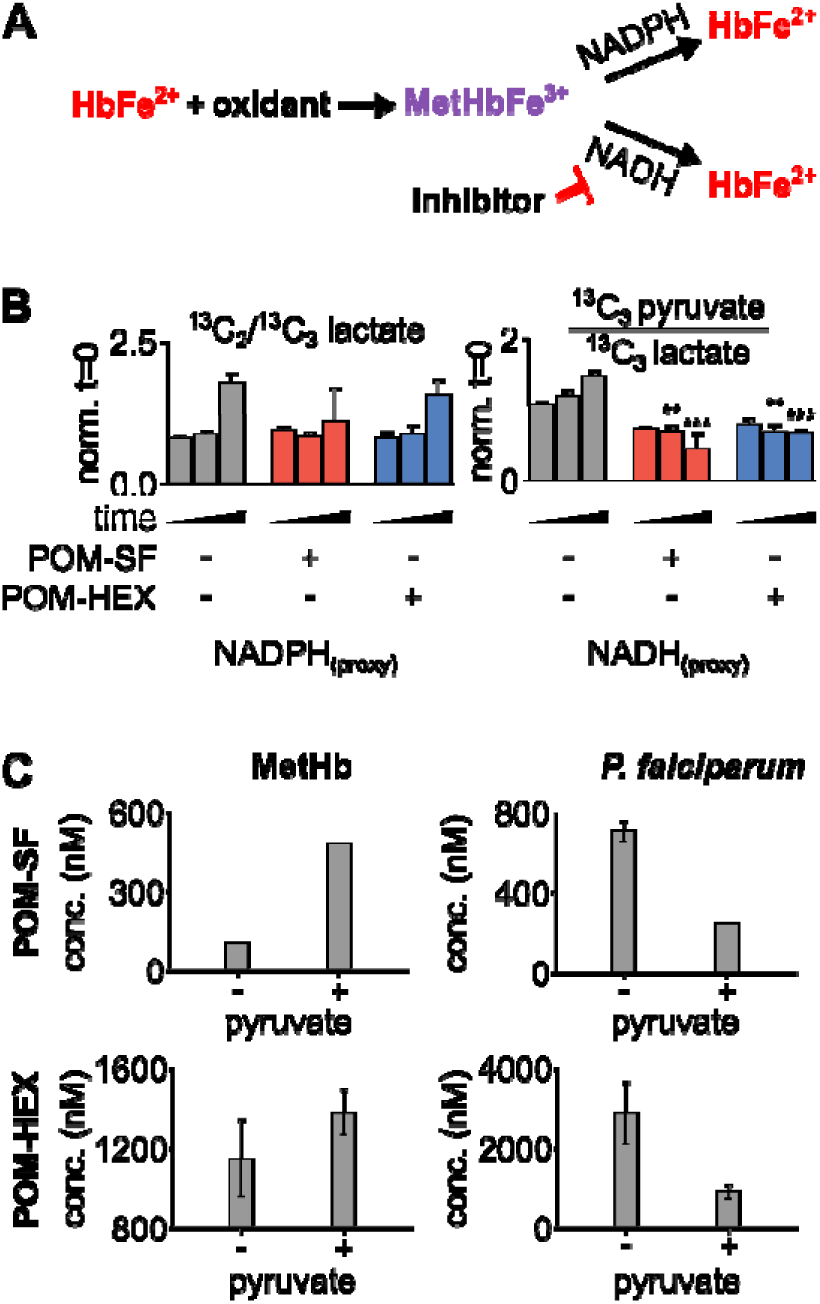
Loss in reductive capacity induces erythrotoxic oxidative damage and antiparasitic effects. (A) Schematic showing two predominate mechanisms of methemoglobin reduction with (B) glucose labeling revealing a collapse in cellular redox potential with pyruvate M+3/lactate M+3 as a proxy for the cellular NAD+/NADH ratio, coupled without an increase in flux through the pentose phosphate pathway as shown via lactate M+2/lactate M+3 (n=3). (C) On the left, half-maximal inhibitory concentrations (EC_50_) for methemoglobin formation in cultured human erythrocytes, with and without 1 mM supplemental pyruvate. On the right, EC_50_ of *P. falciparum* growth, with and without 1 mM supplemental pyruvate. Representative dose-response curves for can be found in Figure 4 and Supplemental Figure 2. All values are determined from three biological replicates, some error bars are below limit of visualization.

**Figure 4.**
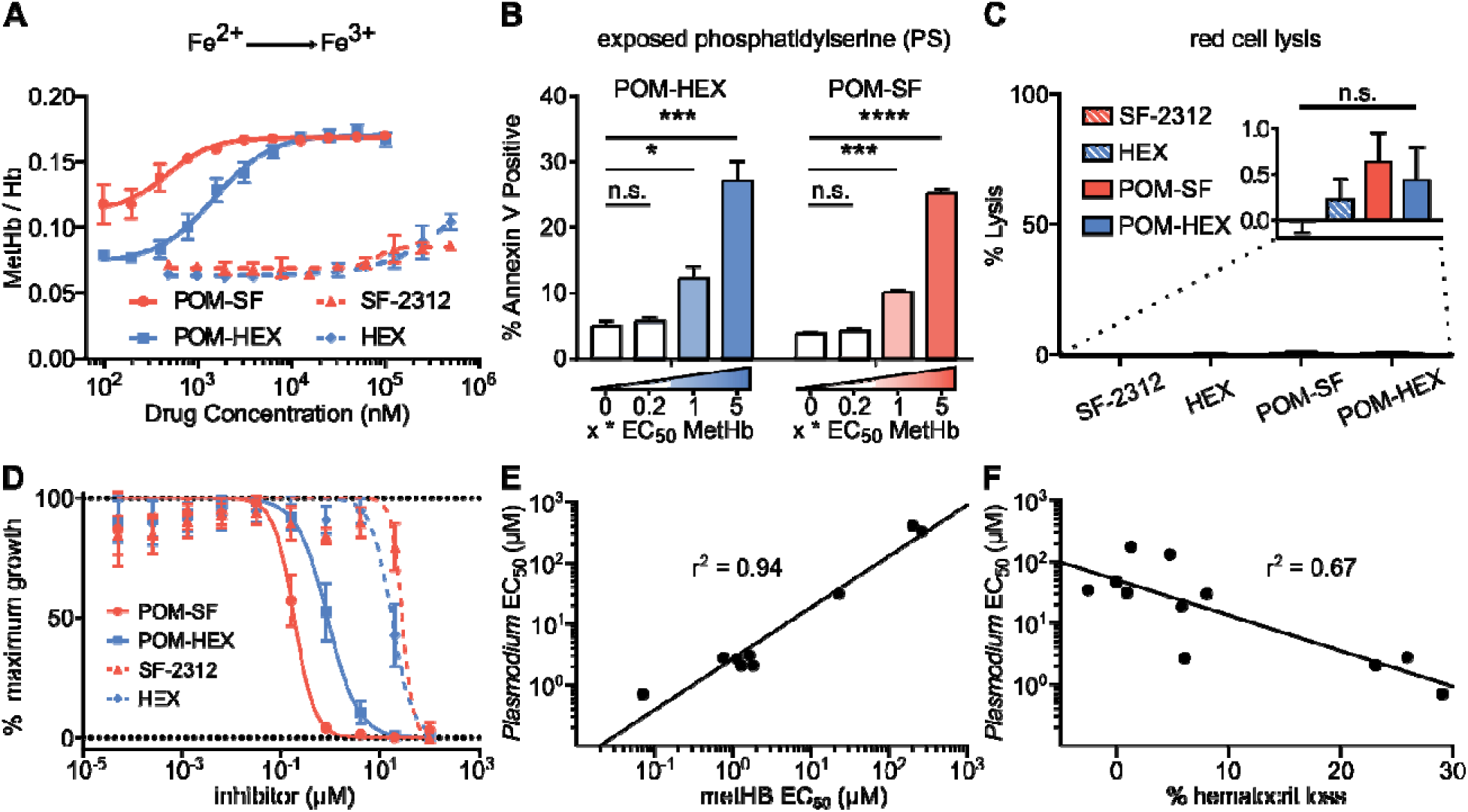
Oxidative damage and erythrocyte senescence upon glycolytic inhibition without hemolysis offers potent antimalarial effects in vitro. (A) Dose-dependent increase in the proportion of oxidized (ferric) heme compared to reduce (ferrous) heme following treatment with indicated enolase inhibitors. Half-maximal effective concentrations are as indicated in Supplemental Table 7 (n=3). (B) Dose-dependent increase in a marker of erythrocyte senescence (exposed phosphatidylserine), characterized via annexin V staining. Concentrations tested are in relation to the EC_50_ of MetHb formation for each compound respectively (n=3). (C) Hemoglobin absorption (normalized to 100% detergent-mediated lysis) in freshly cultured human erythrocytes with and without indicated inhibitors (>100μM for 3 days) (n=3). (D) EC_50_s were performed against *P*. *falciparum* strain 3D7. Displayed are representative curves for a subset of the most potent prodrugs and their respective parent compounds. EC_50_s against parasite growth for all tested compounds are displayed in Supplemental Table 7. (E-F) A high correlation between parasite killing and host toxicity indicates shared mechanism. A linear regression was performed on all compounds for which an EC_50_ for (E) methemoglobin or (F) in vivo mouse hematocrit loss and parasite growth inhibition could be determined (GraphPad Prism). (A, D) The respective EC_50_s are calculated using non-linear regression from each of the independent biological replicates (GraphPad Prism). Hb = hemoglobin, met-Hb = methemoglobin.

Cellular insults, including oxidative damage, cause erythrocytes to be marked for removal from circulation. Senescent erythrocytes are characterized by a loss of membrane asymmetry that results in extracellular exposure of phosphatidylserine (PS), detected by annexin-V staining (31–33). While normal cells would respond by promoting PS internalization by ATP-dependent flippases, inhibition of glycolysis is expected to disrupt such repair. As with methemoglobin formation, we find that glycolytic inhibition of human erythrocytes leads to a dose-dependent increase in annexin V positivity (Figure 4B). Together, these results establish that glycolysis is essential to erythrocyte homeostasis. In addition, these studies reveal a likely mechanism by which genetic erythrocyte enzymopathies that impair central carbon metabolism reduce the circulatory half-life of erythrocytes (34–36).

### Acute glycolytic inhibition does not directly cause erythrocyte lysis

Our data support a mechanism by which glycolytic inhibition marks erythrocytes for macrophage-mediated destruction in vivo. In parallel, we evaluated whether acute glycolytic inhibition also leads directly to hemolysis. To our surprise, continuous exposure to high concentrations of enolase inhibitors does not lead to erythrocyte lysis in static culture (Figure 4C). These results indicate that senescence and premature splenic clearance and destruction, rather than direct hemolysis, are the most likely mechanism for the decreased circulating half-life and anemia associated with interrupted glycolysis in erythrocytes in vivo.

### Antimalarial effects of acute inhibition of erythrocyte glycolysis

*Plasmodium* spp. induce host oxidative stress upon erythrocyte infection (37–39). Therefore, inhibitors that impair erythrocyte reductive capacity may result in malaria protection. Erythrocyte-targeted antimalarials would be highly desirable as “resistance-proof” antiparasitics (40–42). We therefore evaluated whether acute inhibition of erythrocyte glycolysis was directly antiparasitic to cultured *P. falciparum* (Figure 4D; Table 1). Treatment of asexual *P. falciparum* with phosphonates (SF-2312 and HEX) directly reduces parasite growth, with half maximal effective concentrations (EC_50_s) > 15μM. Lipophilic ester pro-drugs, which likely exhibit improved cellular penetration (43–45), exhibit 20- to 160-fold increased antimalarial potencies, compared to their cognate phosphonates (Figure 4D; Table 1). The most potent of these, POM-SF (EC_50_ = 250±10nM), exhibits an in vitro potency in the submicromolar range as compared to other antimalarial compounds under development, such as the dihydroorotate dehydrogenase inhibitor DSM265 (EC_50_ = 53nM) and the recently FDA-approved tafenoquine (EC_50_ = 436nM) (46, 47).

**Table 1.**
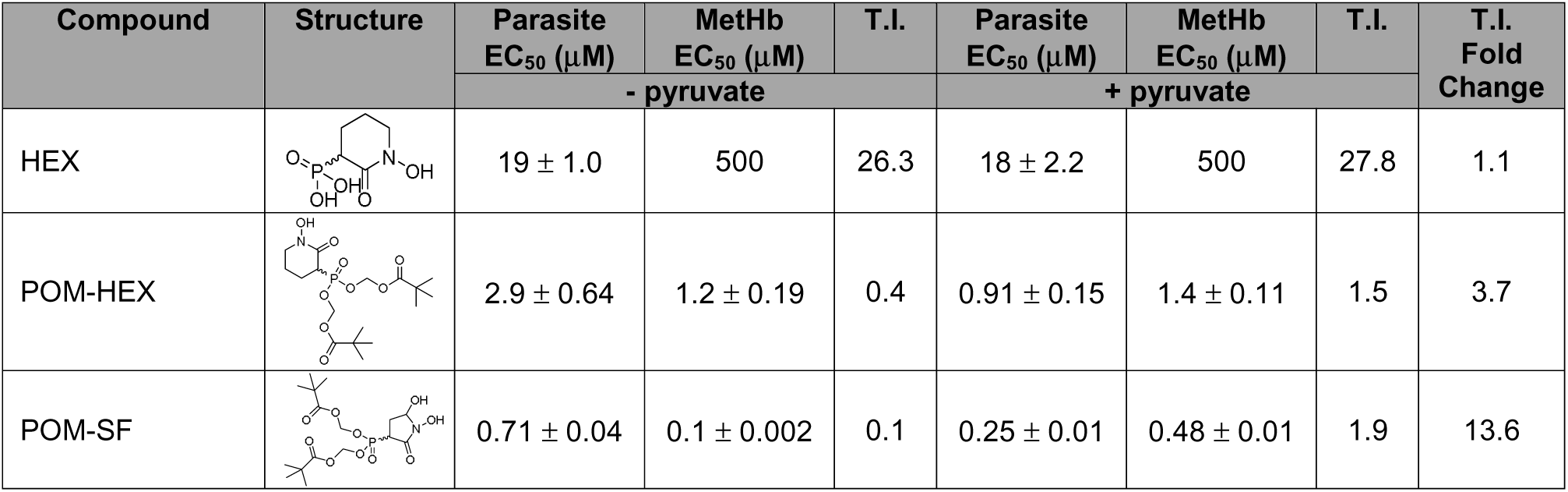
Table of parasite EC_50_s and methemoglobin EC_50_s for all tested compounds and their structures. Displayed are the means ± s.e.m from three or more independent biological replicates in the presence or absence of 1mM pyruvate. EC_50_s are displayed at the highest concentration tested (500 μM) for instances where regression calculations cannot be resolved due to full inhibition extending beyond the highest doses tested. A therapeutic index (T.I.) was calculated as follows: [EC_50_ for MetHb formation] / [EC_50_ for parasite growth inhibition]. The change in T.I. was determined as follows: [T.I. without pyruvate] / [T.I. with pyruvate].

| Compound | Structure | Parasite<br>EC<sub>50</sub> (μM) | MetHb<br>EC<sub>50</sub> (μM) | T.I. | Parasite<br>EC<sub>50</sub> (μM) | MetHb<br>EC<sub>50</sub> (μM) | T.I. | T.I.<br>Fold<br>Change |
| --- | --- | --- | --- | --- | --- | --- | --- | --- |
|  |  | - pyruvate |  |  | + pyruvate |  |  |  |
| HEX |  | 19 ± 1.0 | 500 | 26.3 | 18 ± 2.2 | 500 | 27.8 | 1.1 |
| POM-HEX |  | 2.9 ± 0.64 | 1.2 ± 0.19 | 0.4 | 0.91 ± 0.15 | 1.4 ± 0.11 | 1.5 | 3.7 |
| POM-SF |  | 0.71 ± 0.04 | 0.1 ± 0.002 | 0.1 | 0.25 ± 0.01 | 0.48 ± 0.01 | 1.9 | 13.6 |

Related phosphonate and pro-drug enolase inhibitors were evaluated for their antimalarial potency and their erythrotoxicity. We find that the antimalarial potency of each compound was highly correlated (r^2^=0.94) to its capacity to disrupt erythrocyte redox balance (methemoglobin EC_50_) (Figure 4E). This strict correlation indicates that both effects are likely mediated through a single common target, host erythrocyte enolase. In addition, the degree of inhibitor-induced anemia in vivo in mice also correlates with the antimalarial potency in vitro (r^2^=0.67), suggesting that both anemia and antimalarial potency are mediated by a disruption in host cell redox balance (Figure 4F).

### POM-SF and POM-HEX are effective against multi-drug resistant *P. falciparum*

We further evaluated the antimalarial activity of POM-SF and POM-HEX against parasites with reduced sensitivity to multiple classes of antimalarials, including chloroquine, sulfadoxine-pyrimethamine, mefloquine, and artemisinin. As expected, when targeting erythrocyte biology, multi-drug-resistant parasite lines remain sensitive to both POM-HEX and POM-DF, although there is a modest decrease in POM-SF sensitivity (Supplemental Table 8).

### Chemical complementation with pyruvate improves antiparasitic selectivity

To confirm the mechanism of enolase inhibitor-mediated erythrotoxicity, we evaluated strategies to rescue the erythrotoxicity of enolase inhibition. Pyruvate is a metabolic product immediately downstream of enolase and has been successfully and safely used in humans as a clinical antioxidant, including as a cardio- and neuro-protectant (48–50). We find that exogenous pyruvate ameliorates methemoglobin formation caused by glycolytic inhibition of erythrocytes. In contrast, we find pyruvate potentiates parasite growth inhibition (Figure 3). As a result, pyruvate supplementation markedly improves the therapeutic index of glycolytic inhibitors (Table 1).

### Parasite-selective enolase inhibition ameliorates cerebral malaria

Previous studies indicate that the glucose mimetic 2-deoxyglucose (2-DG) does not reduce parasitemia in acute malaria infection, but yet improves survival in a murine model of cerebral malaria (51, 52). We took advantage of the fact that 2-DG inhibits glycolysis non-specifically in all cell types, while enolase inhibitors (HEX and POM-HEX) specifically target erythrocytes, to evaluate the role of erythrocyte glycolysis in outcomes of malaria infection (Figure 5A-C). As previously observed, 2-DG treatment does not reduce parasitemia; indeed, 2-DG causes a moderate increase in parasite burden [d4, control parasitemia of 7.8±3.3% versus 13.3±3.9% in 2-DG-treated animals, adj. p-value (Bonferroni method) = 0.011] (51, 52). In our hands, 2-DG did not improve the clinical score or overall survival due to cerebral malaria.

**Figure 5.**
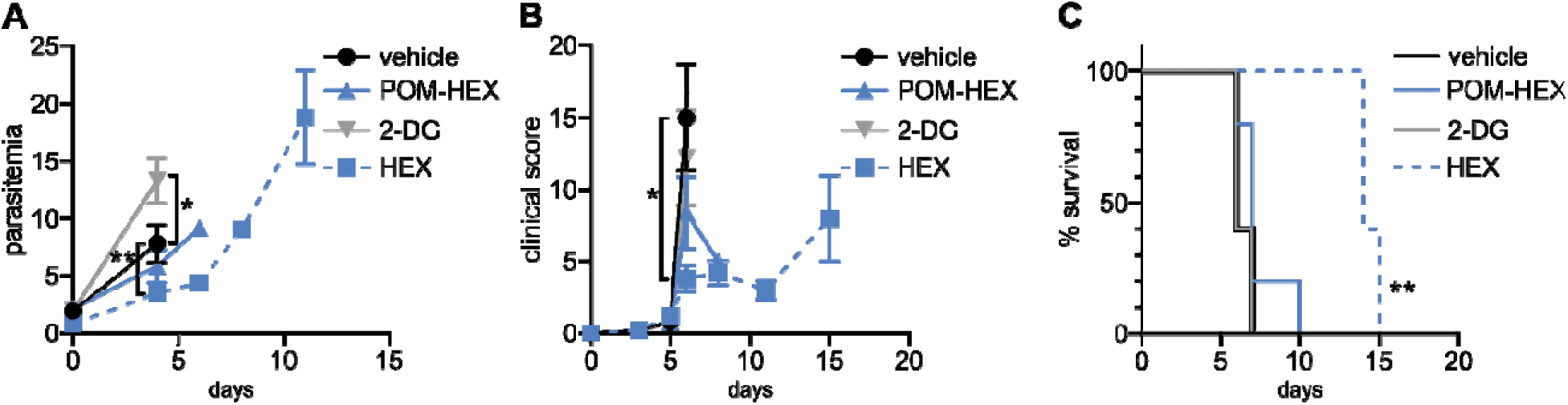
Positive clinical impacts on murine severe malaria following glycolytic inhibition despite high parasitemias. 2-deoxyglucose (2DG) has pleiotropic effects beyond glycolytic inhibition and effects all cell types, while POM-HEX and HEX are specific to ENO2 isozyme expressing cell types like erythrocytes. (A-C) Swiss Webster mice (n=5) were infected with *P. berghei* ANKA strain parasites expressing a luciferase reporter and treated daily from day 2 onwards with either vehicle, 2DG (200 mg/kg), or enolase inhibitors POM-HEX (30 mg/kg), or HEX (100 mg/kg). (A) Mice were monitored for parasitemia via blood smear Giemsa staining on indicated days, * = p ≤ 0.05, ** = p ≤ 0.01. (B) Mice were also assessed for clinical signs of cerebral malaria as determined by signs of neurological symptoms such as paralysis, deviation of the head, ataxia, convulsions, and coma on indicated days, * = p ≤ 0.05. (C) Survival was determined over 15 days, two mice remained alive in the Hex treated group on day 15 but were sacrificed due to high parasitemia concomitant with elevated clinical score as depicted by Kaplan-Meier survival curve, ** = p ≤ 0.01.

Our in vitro studies predicted that erythrocyte toxicity would limit the antimalarial efficacy of the prodrug POM-HEX, which penetrates cells easily, leading to comparable erythrotoxic and antimalarial potencies (Table 1). In contrast, *Plasmodium-*infected erythrocytes actively transport charged molecules. For this reason, the phosphonate parent compound, HEX, is substantially less toxic to uninfected erythrocytes but retains antiparasitic activity (Table 1), predicting a wider therapeutic index in vivo. Indeed, in murine *Plasmodium berghei* infection, we find that HEX treatment reduced overall parasite burden [d4, control parasitemia 7.8±3.3% versus 3.5±1.3% in HEX-treated animals; adj. p-value (Bonferroni method) = 0.009], improved clinical scores [d6, vehicle 15.0±7.3 versus 3.8±2.0 in HEX-treated animals; adj. p-value (Bonferroni method) = 0.039], and significantly improved survival [14 days, compared to 6 days for vehicle-treated animals; adj. p-value (Bonferroni method) = 0.0069 (Mantel-Cox)]. In contrast, POM-HEX, which is non-selectively toxic to both uninfected and infected erythrocytes, was no more effective than 2-DG. These data support the use of in vitro comparisons of erythrotoxicity versus antiparasitic potencies in order to predict in vivo preclinical efficacy. We find that the in vitro therapeutic index better predicts in vivo success compared to in vitro antimalarial potency alone.

### Non-human primates tolerate high doses of HEX with longer half life

Because HEX treatment improves survival in mice with cerebral malaria, we sought to investigate its pharmacodynamic and pharmacokinetic properties in non-human primates as a model for human application. We determined the serum half-life of HEX in cynomolgus monkeys and mice following sub-cutaneous injections. Although mice were treated with higher doses [400mg/kg], HEX-treated non-human primates [100mg/kg] display a higher serum concentration throughout the experiment and a markedly prolonged serum half-life [mice, 6 minutes vs. non-human primates, 1 hour] (Figure 6). Compellingly, we find that the serum concentrations of HEX remain above the in vitro EC_50_ concentrations against *Plasmodium* for at least four hours. Improved drug exposure in non-human primates bodes well for enhanced antimalarial efficacy in humans.

**Figure 6.**
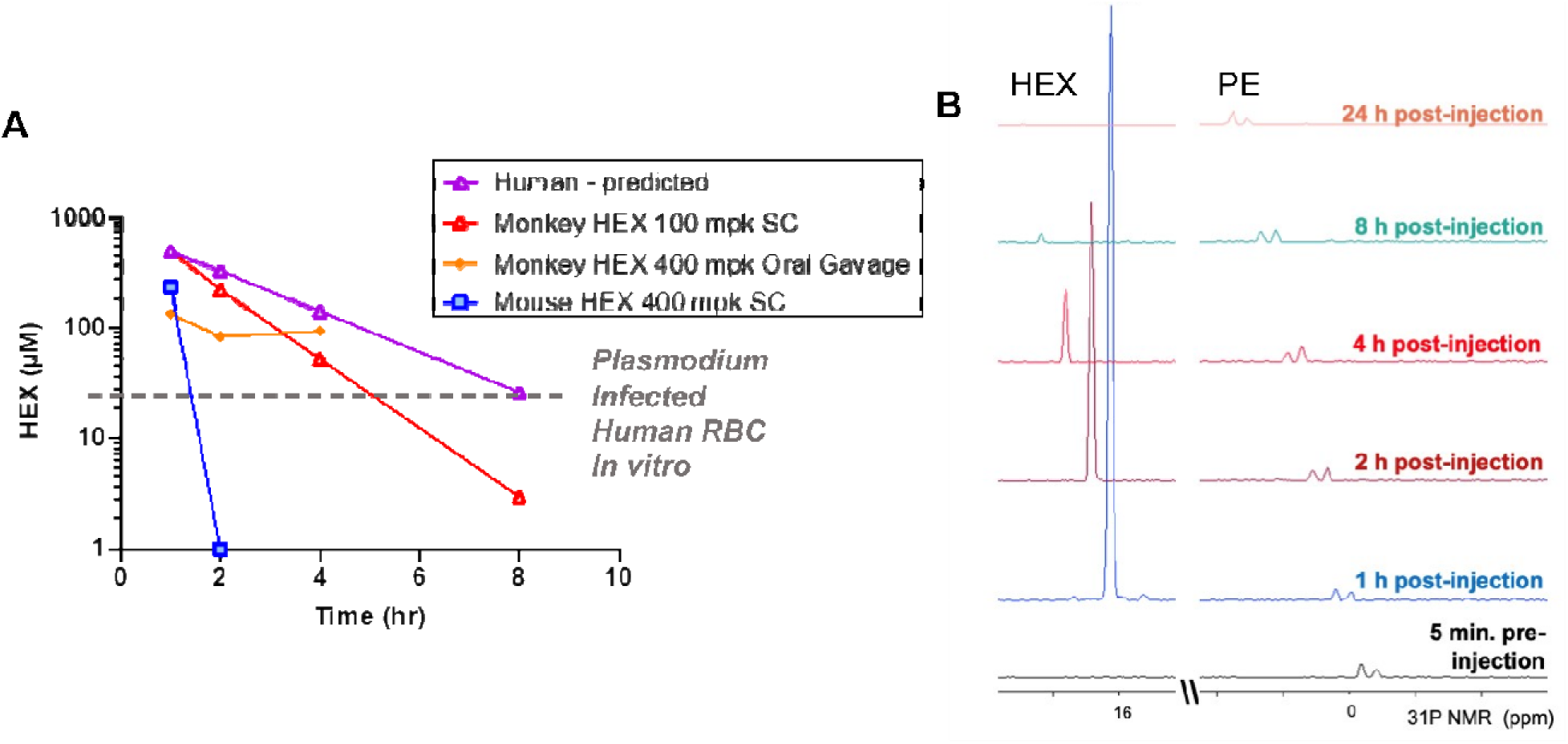
High plasma exposures of HEX are tolerated in Non-human Primates. (A) HEX concentrations in plasma were measured by 31P-1H HSQC (NS = 128) with a detection limit of ∼1 µM. A 400 mg/kg dose SC in mice yielded ∼250 µM HEX 1hr after injection, becoming undetectable at the 2 hr time point. In contrast, a dose of just 100 mpk yielded plasma concentrations of >500 µM in cynomolgus monkeys at 1hr, which remained significant for several hours thereafter. Approximate IC50 concentrations for RBC infected with *Plasmodium* shown as a grey trace, indicating that in mice, sufficient drug concentrations are for killing *Plasmodium* infected RBC are only maintained shortly after injection. The half-lives for HEX in monkey is around 1 hr, whilst in mice it is ∼6 minutes, well in line with values obtained for fosmidomycin. Purple trace, predicted PK of HEX in human at a 100 mpk S.C. dose, based on monkey/human fosmidomycin comparison. The yellow trace, 400 mpk HEX administrated by oral gavage in monkey, indicates HEX is orally bioavailable. B, NMR 31P-1H HSQC 1D read outs at different time points post S.C. HEX (16 ppm chemical shift) and endogenous phosphate esters (PE), indicated.

**Figure 7.**
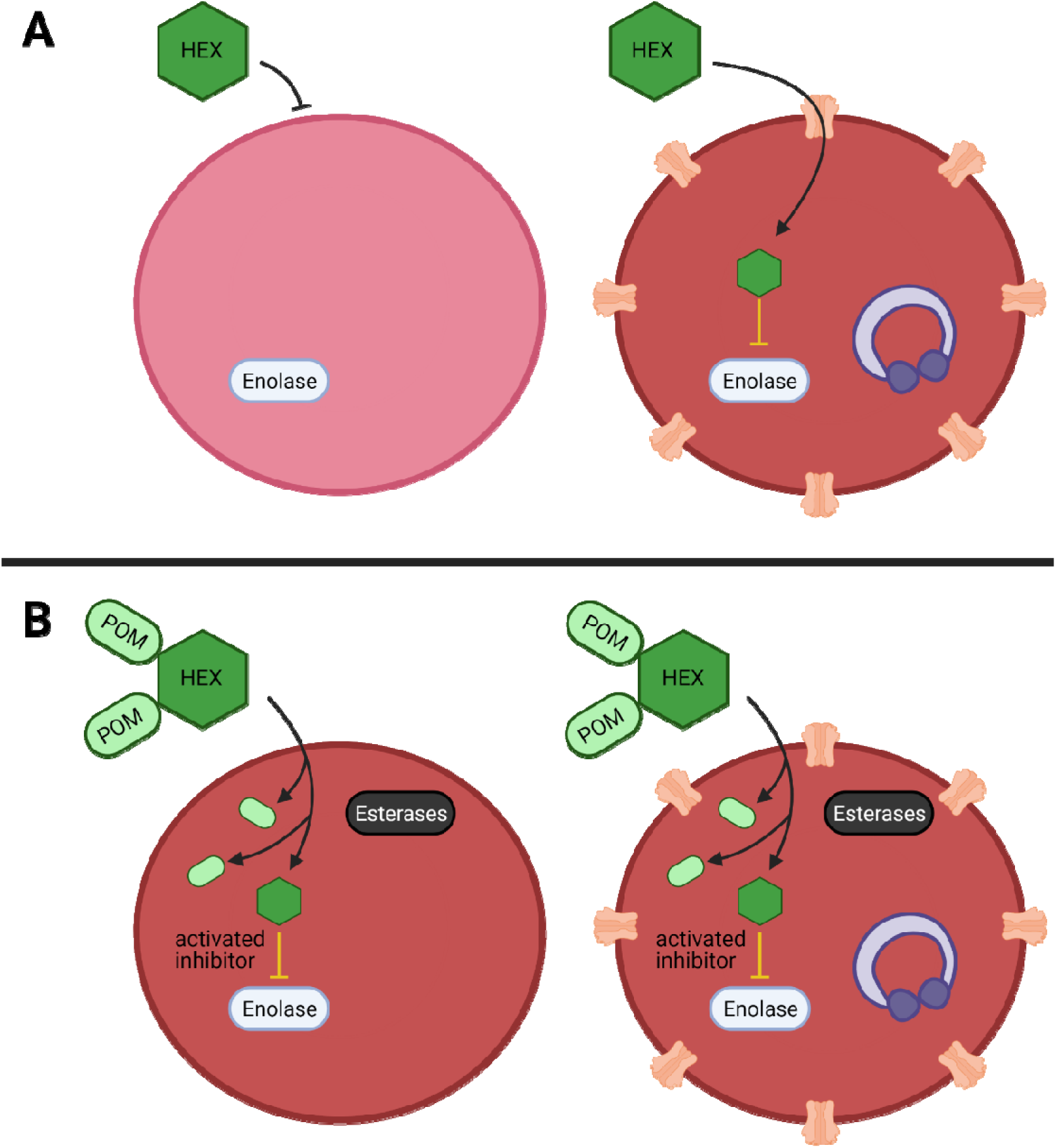
Mechanistic model of anti-parasitic selectivity of a host target. (A-B) Darker red indicates increased metHB formation, infected red cells are indicated by purple ring-stage parasites, erythrocyte membrane channels are plasmodium derived promiscuous permeability pathways, prodrug processing is indicated by cytosolic esterases (black) (A) The enolase inhibitor HEX exhibits low potency but is selectively targeted to host infected red blood cells to generate a therapeutic index. (B) The prodrug cognate POM-HEX increases potency but eliminates any therapeutic window. Created with BioRender.com.

## DISCUSSION

As the central backbone of cellular metabolism, glycolysis controls energy production, redox balance, and biosynthesis. Antiglycolytic inhibitors—so-called “starvation mimetics”—are under increasing consideration as potential therapies for conditions including obesity, cancer, aging, autoimmunity, and infectious diseases (53–57). Most recently, it has been reported that host glycolytic inhibition markedly reduced replication of SARS-CoV-2 (58). While glycolytic disruption is generally well tolerated, glycolysis is critical to metabolic homeostasis in erythrocytes, which exclusively rely on the Embden-Meyerhof-Parnas pathway for ATP generation. As such, anemia is a key dose-limiting side effect and presents a major threat to otherwise promising antiglycolytic therapies (27, 59). Our work provides a mechanistic model through which to understand acute inhibition of erythrocyte glycolysis.

Inhibition of erythrocyte glycolysis is a particularly compelling target for antimalarial therapies. Not only are the symptomatic asexual stages of *Plasmodium* spp. housed within the red cell niche, but long-standing evidence suggests that the outcome of clinical malaria is intertwined with red cell metabolism. Most notably, heritable erythrocyte disorders that appear to reduce malaria severity, including enzymopathies such as G6PDd and PKd, dysregulate glycolytic function and antioxidant capacity (1, 2). Because hemoglobin binds and competes for band 3 sequestration of glycolytic enzymes, malaria-protective hemoglobinopathies such as sickle cell trait also disrupt glycolytic regulation and sensitize erythrocytes to oxidative stress (2). These natural genetic experiments suggest that modulating erythrocyte metabolism successfully improves malaria outcomes, and underscore that long-standing disruptions in erythrocyte metabolic homeostasis can be reasonably well tolerated. Although several antimalarials are contraindicated by glycolytic enzymopathies and novel anti-glycolytic therapies such the reported enolase inhibitors should consider this in future development. As *Plasmodium* spp. have a remarkable capacity for drug resistance, targeting host factors for antimalarial discovery is highly attractive to slow emergence of resistance.

However, our data demonstrate that non-discriminately disrupting glycolysis in infected and uninfected erythrocytes neither drives antiparasitic activity nor improves the outcome of acute malaria infection. Potent enolase inhibition does not forestall a lethal parasite burden (POM-SF and POM-HEX) and does not prevent cerebral malaria (POM-HEX). Rather, our non-prodrugged phosphonates (e.g., HEX) reduce parasite burden, extend survival, and decrease clinical symptoms of malaria. A possible explanation for the discrepancy in antimalarial efficacy of POM-HEX in vitro vs in vivo is that POM-HEX is rapidly converted to Hemi-POM-HEX in mouse plasma, such that effective plasma exposure to POM-HEX itself is quite low (27). The high carboxylesterase activity in mouse (compared to human) plasma has been a recurrent complication for modeling the effects of POM-containing pro-drugs and suggests that the performance of POM-HEX in mice likely underestimates its efficacy in humans (60). The discrepancy between mouse pharmacokinetics and primates is further highlighted in our study, in which the half-life of POM-HEX in primates is 10-fold longer than that in mice. This observation is combined with the likelihood that the reticulocyte tropism of the *P. berghei* model reduces enolase inhibitor efficacy due to increased metabolic plasticity of reticulocytes. The deadliest form of human malaria *P. falciparum* preferentially infects mature erythrocytes.

Because our compounds exhibit a tight correlation between antiparasitic and erythrotoxic potencies, both phenotypes are likely due to inhibition of a single common cellular target, human enolase. Our studies cannot unequivocally rule out other potential secondary targets, including the *P. falciparum* enolase, which itself is likely essential for intraerythrocytic parasite development. Because *Plasmodium falciparum* is an obligate intracellular parasite, it is challenging to evaluate cellular parasite enolase inhibition in isolation. In support of human enolase as the primary antimalarial target of these compounds, however, we also find little or no antiparasitic selectivity for most inhibitors in this series. Nonetheless, multiple-target agents that simultaneously inhibit the host and parasite enzymes would be expected to have durable, resistance-resistant antiparasitic activity (61).

Among our enolase inhibitors, two compounds are notably selective for antiparasitic activity over erythrocyte toxicity, the unmodified parent phosphonates, specifically HEX and DeoxySF-2312. During intraerythrocytic development, *Plasmodium* spp. parasites establish new permeability pathways on the host erythrocyte membrane and are thus relatively permissive to small anions, specifically including phosphonates (62–64). Compared to uninfected red cells that lack these transporters, *Plasmodium*-infected red cells are thus more susceptible to enolase inhibition with charged phosphonate inhibitors. Therefore, we suggest that small molecule inhibitors designed to take advantage of the increased permeability of infected erythrocytes to specifically target glycolysis in *Plasmodium*-infected—but not uninfected—erythrocytes may represent a novel, successful, “resistance-proof” antimalarial strategy. Rapid compound clearance likely ameliorates the therapeutic effect of phosphonates such as HEX in mice, and our studies suggest that primates may represent a better pre-clinical model due to increased circulating half-life.

Our studies also establish a mechanistic model for anemia caused by acute antiglycolytic therapeutics through rapid premature erythrocyte senescence, rather than direct hemolysis. Targeted enolase inhibitor treatment of erythrocytes leads to a metabolic signature characterized by a collapse in glutathione homeostasis. The antioxidant capacity of erythrocytes is dependent on energy production, as erythrocytes, lacking mitochondria, are uniquely dependent on aerobic glycolysis. Enolase inhibition reduces the net ATP per glucose molecule by half, thus hampering production of the energy-expensive cellular antioxidant, glutathione.

The production of this tripeptide (glutamate, cysteine, and glycine) requires ATP at each assembly step. Thus, enolase inhibition globally disturbs amino acid metabolism and leads to a state of energy depletion and the onset of oxidative stress. The depletion of glutathione likely results in an accumulation of oxidative insults over time that results in the metHb formation observed.

Depletion of reduced glutathione establishes the molecular mechanism of methemoglobin formation and senescence in inhibitor-treated erythrocytes. Lacking organelles, erythrocytes undergo a distinct cell death known as eryptosis. Like apoptosis, eryptosis causes cell shrinkage, membrane blebbing, protease activation, and loss of membrane asymmetry. Changing membrane phospholipids (e.g., phosphatidylserine) serves as a removal signal for circulating macrophages and splenic clearance. Our investigations thus explain the features of anemia observed in enolase-inhibitor-treated mice, who develop profound splenomegaly but not circulating extracellular hemoglobin (as would be expected by hemolysis).

As antiglycolytic therapies proceed through development, strategies to ameliorate erythrotoxicity will be required. We find that exogenous pyruvate supplementation restores reductive capacity to inhibitor-treated erythrocytes. This agrees with previous findings that pyruvate alone maintains erythrocyte ATP levels and 2,3-diphosphoglycerate (2,3-DPG) levels during long-term storage (65, 66). In contrast, pyruvate potentiates the antiparasitic effect of enolase inhibition, perhaps through a feed-back inhibition of parasite glycolysis. These data suggest a profound biological difference in glycolytic regulation between *Plasmodium*-infected and uninfected erythrocytes that may be harnessed for future antimalarial therapeutics. In addition, well-tolerated oral pyruvate supplementation may represent a useful erythroprotective strategy alongside antiglycolytic “starvation-mimetic” therapies—to have one’s cake, and eat it, too (67–69).

## METHODS

### Compounds

Phosphonoacetohydroxamate (Sigma), SF2312 (Chem Space), DeoxySF2312, HEX, POM-HEX, and POM-SF were synthesized per previously published procedures (25, 27). Synthesis of novel SF2312 derivatives and HEX-pro drugs was performed as described (70).

### Washing, storage, and in vitro culture conditions of erythrocytes

Banked blood (Saint Louis Children’s Hospital) and fresh blood [healthy donors] was washed and stored at 50% hematocrit at 4°C up to 1 month past the clinical expiration date and collection date, respectively. Parasite strains provided by the Malaria Research and Reference Reagent Resource Center (MR4) as follows: 3D7 (MRA-102); K1 (MRA-159); D10 (MRA-201); and IPC-5202 (MRA-1240). Unless indicated, erythrocytes were cultured as previously described at 2% hematocrit in complete media (incomplete media supplemented with 27mM sodium bicarbonate, 11mM glucose, 5mM HEPES, 1mM sodium pyruvate, 0.37mM hypoxanthine, 0.01mM thymidine, 10μg/mL gentamicin, and 0.5% Albumax (Thermo-Fisher) under 5% O_2_/5% CO_2_/90% N_2_ atmosphere at 37°C (71).

### *Plasmodium falciparum* growth inhibition assays

Cultures diluted to 1% parasitemia in 100μL wells were subjected to drug dilutions, solvent controls, and untreated uninfected erythrocyte controls in technical duplicates. Following a three-day incubation, DNA content was quantified using Picogreen (Life Technologies), as previously described (72). Half-maximal effective concentrations (EC_50_) were determined from nonlinear regression of three biological replicates (GraphPad Prism).

### Detection and quantification of in vitro erythrocyte hemolysis

Erythrocytes were treated with 100μM of each compound for 72 hours. Supernatants were collected and erythrocytes were resuspended in 100μL complete media containing 0.1% saponin and diluted 1:10. Hemoglobin (Hb) content of fully lysed erythrocytes and supernatants were determined via Abs 404nm. Percent lysis was defined (from four replicates across three separate blood donors) as the ratio of [supernatant Hb content]/[Hb content of fully lysed cultures] X 100.

### Detection and quantification of methemoglobin formation

Methemoglobin (MetHb) was detected by measuring absorbance in the 630-640nm range, which is specific from other hemoglobin subforms (oxyhemoglobin, deoxyhemoglobin, and carboxyhemoglobin). Supernatants were removed and erythrocyte pellets were fully lysed and diluted, as described above. The absorption peak for oxyhemoglobin (575nm) and methemoglobin (630nm) were measured and the EC_50_ of MetHb formation was determined using non-linear regression analysis (GraphPad Prism).

### Determination of erythrocyte membrane asymmetry

Membrane asymmetry was determined by measuring Annexin-V labeling on day three of compound treatment. Erythrocytes washed with 200μL Annexin-V Binding Buffer (140mM sodium chloride, 2.5mM calcium chloride, 10mM HEPES pH 7.4) were briefly spun, supernatants removed, and resuspended in buffer containing Annexin-V conjugated to Alexa-Fluor^TM^ 488 (Invitrogen A13201). Annexin-V positivity was determined using an LSR-II BD FACS analyzer containing a 488nm laser. Eryptotic erythrocytes were compared to a 5μM ionomycin-treated positive control and an untreated negative control.

### Retro-orbital bleeding and hematocrit determination

Mice were anesthetized using isoflurane. Blood was collected using heparinized microhematocrit capillary tubes (Fisherbrand #22-362574) via retro-orbital approach and was sealed with hemato-seal capillary tube sealant (Fisherbrand #02-678). After spinning at 1,500xg for 10 min, hematocrit was calculated by dividing the measured height of red blood cell layer by the total height of the blood.

### Sample collection for untargeted metabolomics

Freshly collected erythrocytes were washed, resuspended to 10% hematocrit in buffer [25mM HEPES (pH 7.4), 120mM NaCl, 5.4mM KCl, 1.8mM CaCl_2_, and 1mM NaH_2_PO_4_], incubated for 1 hour at 37°C, then centrifuged at 2000xg for 5 minutes at 4°C, resuspended in fresh wash buffer to 30-40% hematocrit, and split into 210μL aliquots of packed erythrocytes. Erythrocytes were resuspended in 253μL of RPMI containing 11.9mM D-[1,2,3-^13^C] glucose (Millipore-Sigma) and either treated with POM-SF or POM-HEX at five times the EC_50_s for MetHb formation, or an untreated solvent control with the balance volume to 550μL using washing buffer. All samples were incubated at 37°C shaking at 500 RPM. Triplicate samples of supernatants and packed erythrocytes were collected at 0, 0.5, 1, and 6-hour intervals and snap-frozen in liquid N_2_.

### Sample processing, metabolite extraction, and metabolite detection

RBC and media samples were extracted at 1:10 and 1:20 dilutions, respectively, in 5:3:2 MeOH:MeCN:H2O *v/v/v*, and supernatants analyzed on a Thermo Vanquish UHPLC coupled to a Thermo Q Exactive mass spectrometer, as extensively described (73, 74). The extraction buffer was supplemented with 40uM 3-^13^C lactate to allow absolute quantification of lactate isotopologues, including lactate generated via glycolysis ([1,2,3-^13^C_3_] enriched) vs. the pentose phosphate pathway ([2,3-^13^C_2_] enriched), as described (75). Metabolite assignments, peak area integration and isotopologues distributions were determined using Maven (Princeton University) (73, 74).

### Metabolite data processing and analysis

Data were auto-scaled and normalized to zero-hour time points. Metabolites were removed from the dataset if greater than 50% of the samples contained missing values or the relative standard deviation among replicates exceeded 25%. Additionally, data was filtered using the interquartile range to reduce near-constant values from the experiment by 5%.

A principal component analysis of preprocessed data was plotted with the 95% confidence interval minimum volume enclosing ellipsoid for each respective group (MetaboAnalyst, cluster, plotly). A two-way repeated measures analysis of variance (ANOVA) was performed with multiple comparisons corrected for using a false discovery rate p-value cutoff of 0.05. Metabolite-Pathway analysis was performed using the R package of MetaboAnalyst, as previously described (76). Metabolite-Protein nodal analysis was performed using omicsnet.ca and as previously described (28).

### Determining in vivo efficacy of enolase inhibitors against blood stage parasite burden

Groups of five female Swiss-Webster mice were injected intraperitoneally (i.p.) with 10^3^ luciferase-expressing *P. berghei* ANKA parasites (77). Forty-eight hours post infection, mice were injected i.p. daily for five days with either 200μL of 2% methylcellulose/0.5% Tween80, 40mg/kg chloroquine, 100mg/kg HEX, 30mg/kg POM-HEX, or 5mg/kg POM-SF. Enolase inhibitors dosing were selected as approximately 3-fold lower than the maximum acutely tolerated dose for each compound by i.p. injection. Day 7 post-infection, mice were injected with 150mg/kg D-luciferin potassium salt in PBS (GoldBio) and imaged on an IVIS 100 and parasite burden assessed (Xenogen LivingImage).

### Determining in vivo efficacy of enolase inhibitors against cerebral malaria

The glucose mimetic 2-DG and enolase inhibitors POM-HEX, and HEX were evaluated in the *P. berghei* mouse model of cerebral malarial, as previously described (52). Briefly, in addition to groups of mice dosed as described above for POM-HEX and HEX, 2-DG was administered to an additional group at 200mg/kg twice daily and all were monitored for survival, parasitemia (blood smear Giemsa), and clinical signs of cerebral malaria (paralysis, deviation of the head, ataxia, convulsions, and coma) over 15 days.

### Pharmacodynamics (PD) and pharmacokinetics (PK) studies of HEX in non-human primates

Adult male cynomolgus monkeys (*Macaca fascicularis*) with weight of 2.5-3.5 kg were used for PK and PD studies. To prepare HEX for injection, a stock solution of 150 mg/mL HEX in water with final pH of 7.2-7.4 was prepared. Just prior to injection, this stock was diluted with saline to desired concentration and filtered with 0.22 µm filter. A dose of 100 mg/Kg of HEX was injected subcutaneously in an overnight fasted monkey. Blood samples were collected at the following time points: before injection, 1 hr, 2 hrs, 4 hrs (when food was also given to a monkey), 8 hrs, and 24 hrs after injection. Also, a dose of 400 mg/Kg of HEX was administrated through oral gavage. To compare the PK studies in non-human primates with rodents, a 400 mg/kg dose of HEX was also injected subcutaneously in mice, and blood was collected before, 1 hr, and 2 hrs after injection. HEX concentrations in plasma were measured by using nuclear magnetic resonance (NMR) spectroscopy. Two hundred microliters of plasma were extracted with four hundred microliters of −20□°C precooled methanol and incubated in −20□°C for 45 minutes followed by spinning down in 4□°C at 17,000□×□g for 30□min to separate supernatant. The supernatant then was concentrated by speedvac for 4 hours and dissolved in 470 μL D2O (Sigma Aldrich,151882) and 30 μL D2O with 3%TPS as an internal standard (Sigma Aldrich, 450510) for NMR studies. NMR experiments were performed on Bruker Avance III HD 500 MHz spectrometer equipped with cryoprobe and Bruker 300 MHz with broad band observe probe. For each sample, we obtained one-dimensional (1D) proton (1H) spectrum using zg30 pulse program, 1D 1H spectrum with inverse gated 31-phosphate (31P) decoupling using zgig pulse program and (two-dimensional) 2D 1H-31P heteronuclear single quantum correlation (HSQC) using hsqcetgp pulse program (with scan parameters of 128 scans, gpz2 %=32.40, 31P SW= 40 ppm, O2p=0 ppm, cnst2=22.95 which is adjusted for HEX J2 coupling). NMR spectra were analyzed using 3.1 version of TopSpin. HEX was quantified in the sample with high concentration of the drug (100 mpk, 1 hr after SC injection) by comparing integrals of HEX and TSP (reference) in 1H spectra. To calculate HEX concentration for other time points, we obtained the 1D projections of 2D 1H-31P HSQC spectrum using the “proj” commend in TopSpin. In the projected spectra, the peak at 16 ppm chemical shift appeared only in HEX treated samples which we used to calculate HEX concentration. We took the ratio of integral of 16 ppm peak for samples with unknown concentration of HEX versus one with known concentration.

### Lead Contact and Materials Availability

Further information and requests for resources and reagents should be directed to and will be fulfilled by the Lead Contact, Audrey Odom John.

### Statistics

All statistics represent a minimum of three biological replicates and all analyses were adjusted for multiple comparisons when necessary. Each statistical analysis is described in the respective figure legend and/or methods section. A minimum P-value threshold of p<0.05 was required for statistical significance.

### Study Approval

Experiments and procedures on non-human primates were performed as fee-for-service by Charles River Laboratories with approval of Charles River’s Institutional Animal Care and Use Committee (IACUC). Mice experiments were performed at University of Texas MD Anderson Cancer Center with approval of MD Andersons’s IACUC. The *P*. *berghei* mice experiments were performed as fee-for-service by the Anti-infectives core at New York University with approval of the NYU IACUC. Fresh human blood was acquired under Washington University protocol (IRB ID #: 201012782) with written informed consent.

## ACKNOWLEDGEMENTS

The authors are grateful to Stephen Rodgers and Allan Doctor for helpful discussion. We thank Sudhir Ragavan and Pijus Mandal for technical assistance in chemical synthesis. We thank Dimitra Georgiou for compliance support. This work is supported by the following: A.J. is supported by NIH T32GM007067. A.D. was supported by funds from the Webb-Waring Early Career award 2017 by the Boettcher Foundation. F.L.M. is an Andrew Sabine family fellow and is supported by The Research Scholar award, RSG-15-145-01-CDD from the American Cancer Society and The Young Investigator Award YIA170032 from the National Comprehensive Cancer Network. A.O.J. is supported by NIH/NIAID R01-AI103280, R21-AI123808, and R21-AI130584, and AOJ is an Investigator in the Pathogenesis of Infectious Diseases (PATH) of the Burroughs Wellcome Fund.

## AUTHOR CONTRIBUTIONS

All authors were involved in experimental design and critical evaluation of the data and manuscript. A.J. performed metabolite sample preparation, red cell and parasite toxicity studies, data analysis, experimental design, data interpretation and manuscript preparation. R. C-H. performed metabolomics analysis. J.A.R. and A.D. performed metabolomics data analysis, data interpretation and manuscript preparation. A.O.J. conceived the study, contributed to experimental design, data interpretation, and manuscript preparation. F.L.M. conceived and synthesized enolase inhibitors; V.Y. synthesized inhibitors and critical review of the manuscript Y.L. carried out hematocrit and toxicity studies. Y.B. performed and analyzed PK and PD studies.

---

**Supplemental Figure 1:**
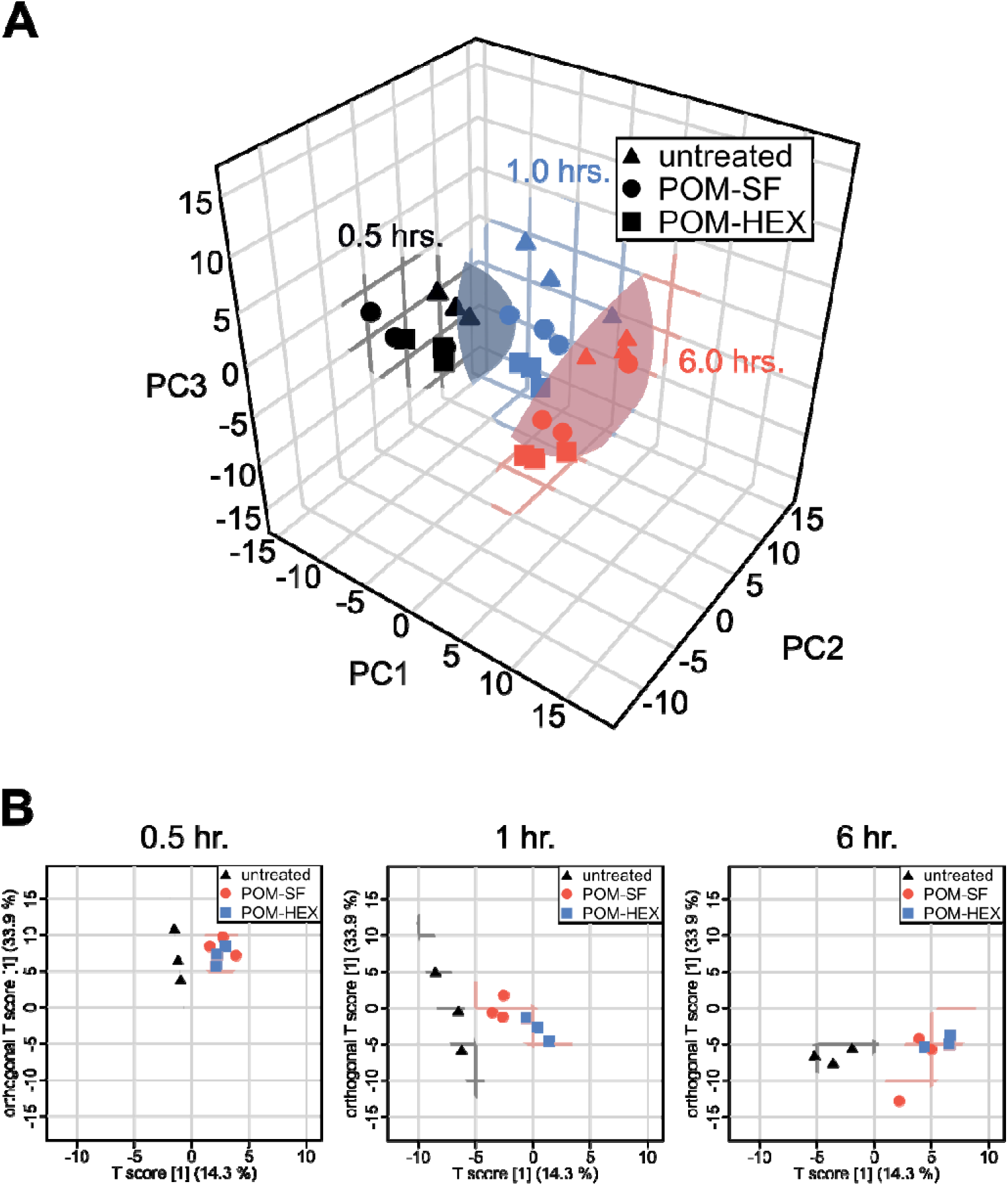
Consistent metabolic signature of glycolytic inhibition. (A) Principle components analysis (PCA) of erythrocytes with and without treatment with enolase inhibitors (POM-SF, 2.4 μM, and POM-HEX, 7 μM) demonstrates distinct separation of samples with respect to time and condition, illustrated by 95% confidence intervals of a minimum volume enclosing ellipsoid for each time point, teal = 0.5 hr., pink = 1 hr., blue = 6 hr. (B) Within each time point there is clear separation of treated samples (circles and squares) from untreated controls (plus symbol) as more clearly illustrated using orthogonal projections to latent structures discriminant analysis (OPLS-DA).

**Supplemental Figure 2:**
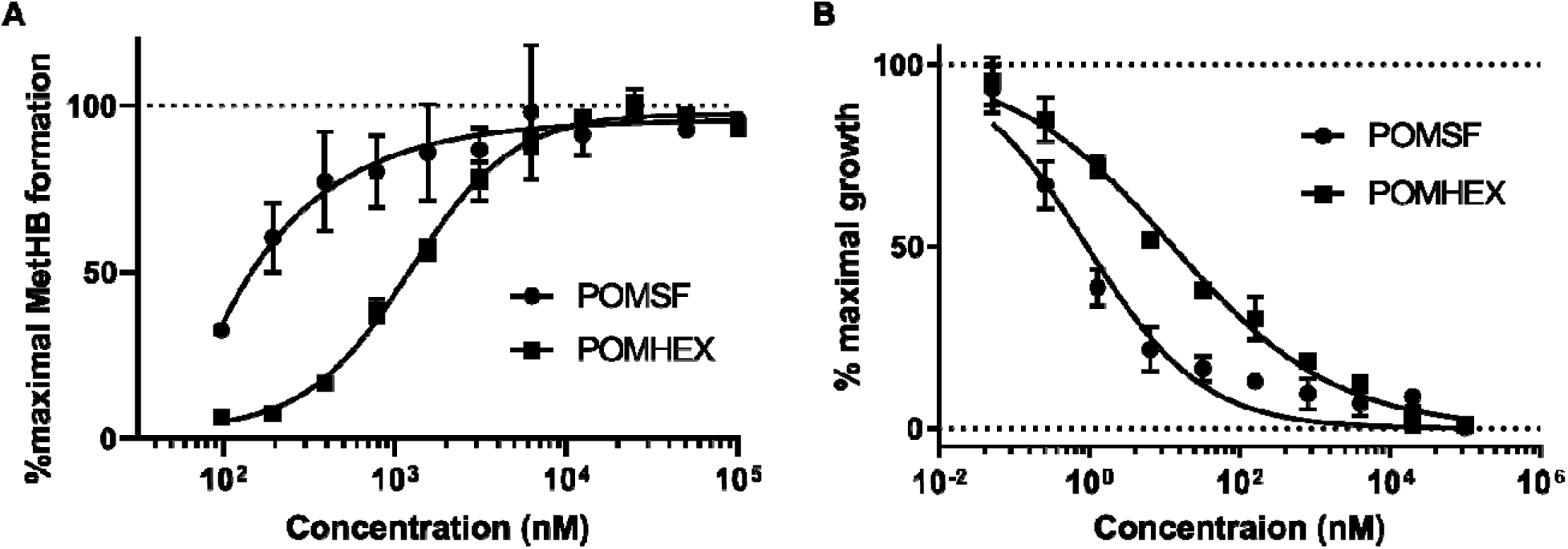
Methemoglobin formation and parasite growth inhibition without pyruvate supplementation. Dose response curves are representative of three independent experimental replicates. Non-linear regression was performed using GraphPad Prism.

## Additional data file

**Supplemental Table 1. Untargeted LC/MS of POM-SF and POM-HEX treated erythrocytes.**

**Supplemental Table 2.** Principle components analysis (PCA). Sample names reflect the sample number followed by the time the sample was collected in units of hours. Results from this table are plotted in Supplemental Figure 1A as a three-dimensional projection in two-dimensional space.

| Sample | Phenotype | Time | PC1 | PC2 | PC3 |
| --- | --- | --- | --- | --- | --- |
| S1T0.5 | Untreated | 0.5 | -11.187 | 4.0977 | -0.50702 |
| S1T1 | Untreated | 1 | -5.2149 | 8.6656 | 4.4176 |
| S1T6 | Untreated | 6 | 6.3029 | 6.6861 | 0.24052 |
| S2T0.5 | Untreated | 0.5 | -3.6239 | -1.1955 | 2.6787 |
| S2T1 | Untreated | 1 | 5.766 | 5.0932 | 3.0898 |
| S2T6 | Untreated | 6 | 7.4986 | 4.2406 | 0.9167 |
| S3T0.5 | Untreated | 0.5 | -6.4935 | 0.68517 | 1.5603 |
| S3T1 | Untreated | 1 | 0.48451 | 3.9959 | 5.1811 |
| S3T6 | Untreated | 6 | 5.4986 | 1.8652 | 0.79194 |
| S4T0.5 | POMSF | 0.5 | -9.1342 | -6.7371 | 3.8178 |
| S4T1 | POMSF | 1 | -1.5747 | 1.2281 | 2.2968 |
| S4T6 | POMSF | 6 | 13.608 | -5.9649 | 7.9595 |
| S5T0.5 | POMSF | 0.5 | -6.6397 | -6.8714 | 2.5662 |
| S5T1 | POMSF | 1 | 1.3937 | 1.8705 | 1.7246 |
| S5T6 | POMSF | 6 | 5.8559 | -4.7354 | -1.2115 |
| S6T0.5 | POMSF | 0.5 | -8.6315 | 1.7468 | -4.032 |
| S6T1 | POMSF | 1 | 0.43739 | 5.597 | -2.188 |
| S6T6 | POMSF | 6 | 3.7712 | 1.4584 | -7.3746 |
| S7T0.5 | POMHEX | 0.5 | -8.2408 | -3.9992 | 0.16622 |
| S7T1 | POMHEX | 1 | 3.209 | -4.12 | 2.6662 |
| S7T6 | POMHEX | 6 | 3.5165 | -3.6213 | -6.4734 |
| S8T0.5 | POMHEX | 0.5 | -5.4074 | -2.5732 | -0.23804 |
| S8T1 | POMHEX | 1 | 1.5453 | 0.018894 | -0.78583 |
| S8T6 | POMHEX | 6 | 4.8119 | -4.0887 | -5.9501 |
| S9T0.5 | POMHEX | 0.5 | -7.4582 | 0.079372 | -4.0643 |
| S9T1 | POMHEX | 1 | 4.9242 | -3.6691 | 0.76423 |
| S9T6 | POMHEX | 6 | 4.983 | 0.24724 | -8.0134 |

**Supplemental Table 3.** Orthogonal projections to latent structures discriminant analysis (OPLS-DA). Sample names reflect the sample number followed by the time the sample was collected in units of hours. Results from this table are plotted in Supplemental Figure 1B for each time point comparing all samples.

| Sample | Phenotype | Time | Score (t1) | OrthoScore (to1) |
| --- | --- | --- | --- | --- |
| S1T0.5 | Untreated | 0.5 | -1.5272 | 10.808 |
| S1T1 | Untreated | 1 | -8.5524 | 5.0246 |
| S1T6 | Untreated | 6 | -5.2479 | -6.5711 |
| S2T0.5 | Untreated | 0.5 | -0.97074 | 3.8803 |
| S2T1 | Untreated | 1 | -6.2673 | -5.7055 |
| S2T6 | Untreated | 6 | -3.6426 | -7.6177 |
| S3T0.5 | Untreated | 0.5 | -1.1952 | 6.5305 |
| S3T1 | Untreated | 1 | -6.5399 | -0.34779 |
| S3T6 | Untreated | 6 | -1.98 | -5.5483 |
| S4T0.5 | POMSF | 0.5 | 2.6948 | 9.6091 |
| S4T1 | POMSF | 1 | -2.6176 | 1.6842 |
| S4T6 | POMSF | 6 | 2.1306 | -12.902 |
| S5T0.5 | POMSF | 0.5 | 3.8568 | 7.0615 |
| S5T1 | POMSF | 1 | -2.6998 | -1.2993 |
| S5T6 | POMSF | 6 | 4.9324 | -5.798 |
| S6T0.5 | POMSF | 0.5 | 1.5729 | 8.3196 |
| S6T1 | POMSF | 1 | -3.5977 | -0.70088 |
| S6T6 | POMSF | 6 | 3.849 | -4.3323 |
| S7T0.5 | POMHEX | 0.5 | 2.9608 | 8.3721 |
| S7T1 | POMHEX | 1 | 0.35493 | -2.7783 |
| S7T6 | POMHEX | 6 | 6.5437 | -3.8174 |
| S8T0.5 | POMHEX | 0.5 | 2.1494 | 5.5285 |
| S8T1 | POMHEX | 1 | -0.63661 | -1.4763 |
| S8T6 | POMHEX | 6 | 6.4988 | -5.0309 |
| S9T0.5 | POMHEX | 0.5 | 2.2136 | 7.2059 |
| S9T1 | POMHEX | 1 | 1.3414 | -4.6182 |
| S9T6 | POMHEX | 6 | 4.3758 | -5.4802 |

**Supplemental Table 4.**
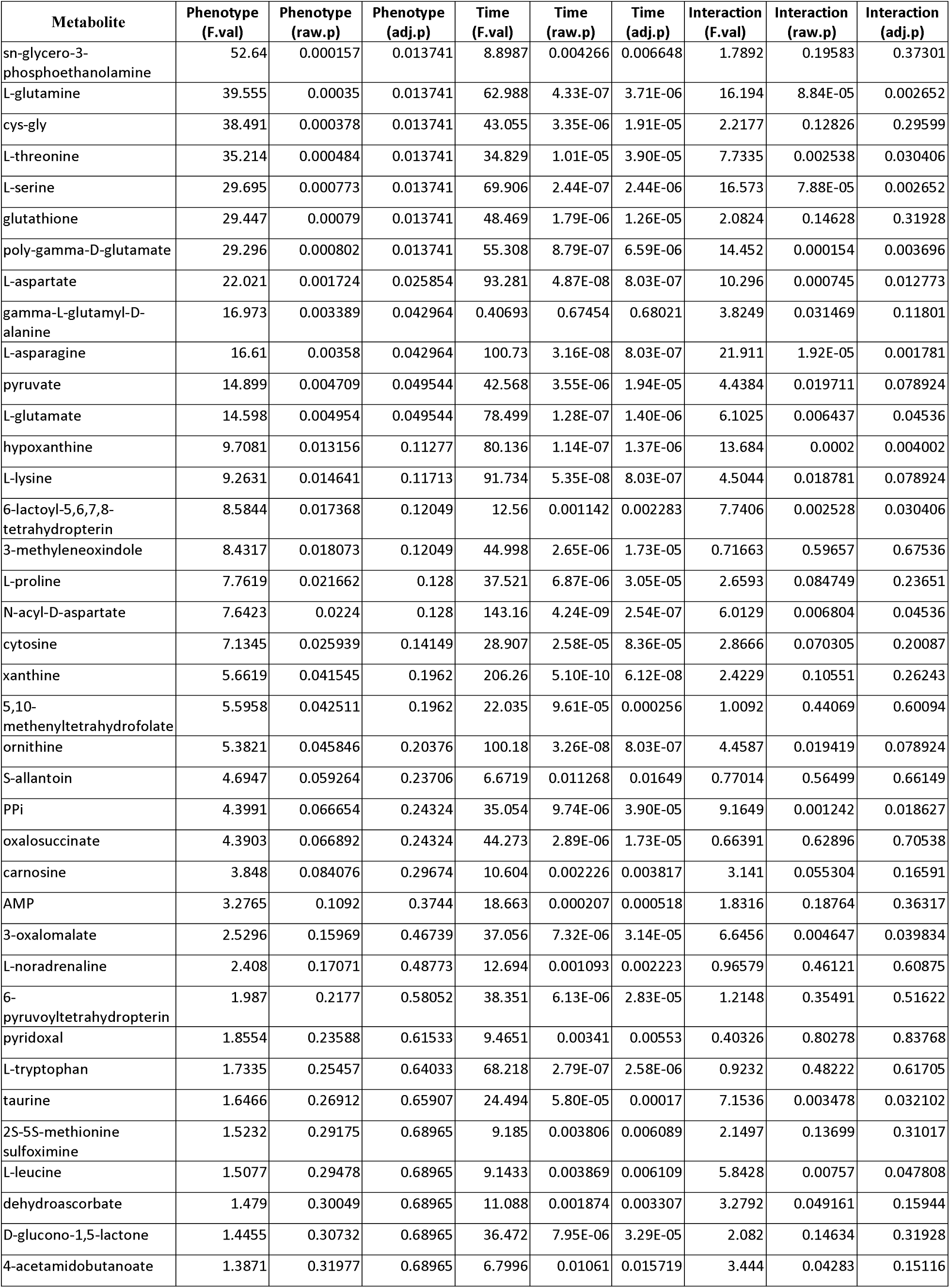
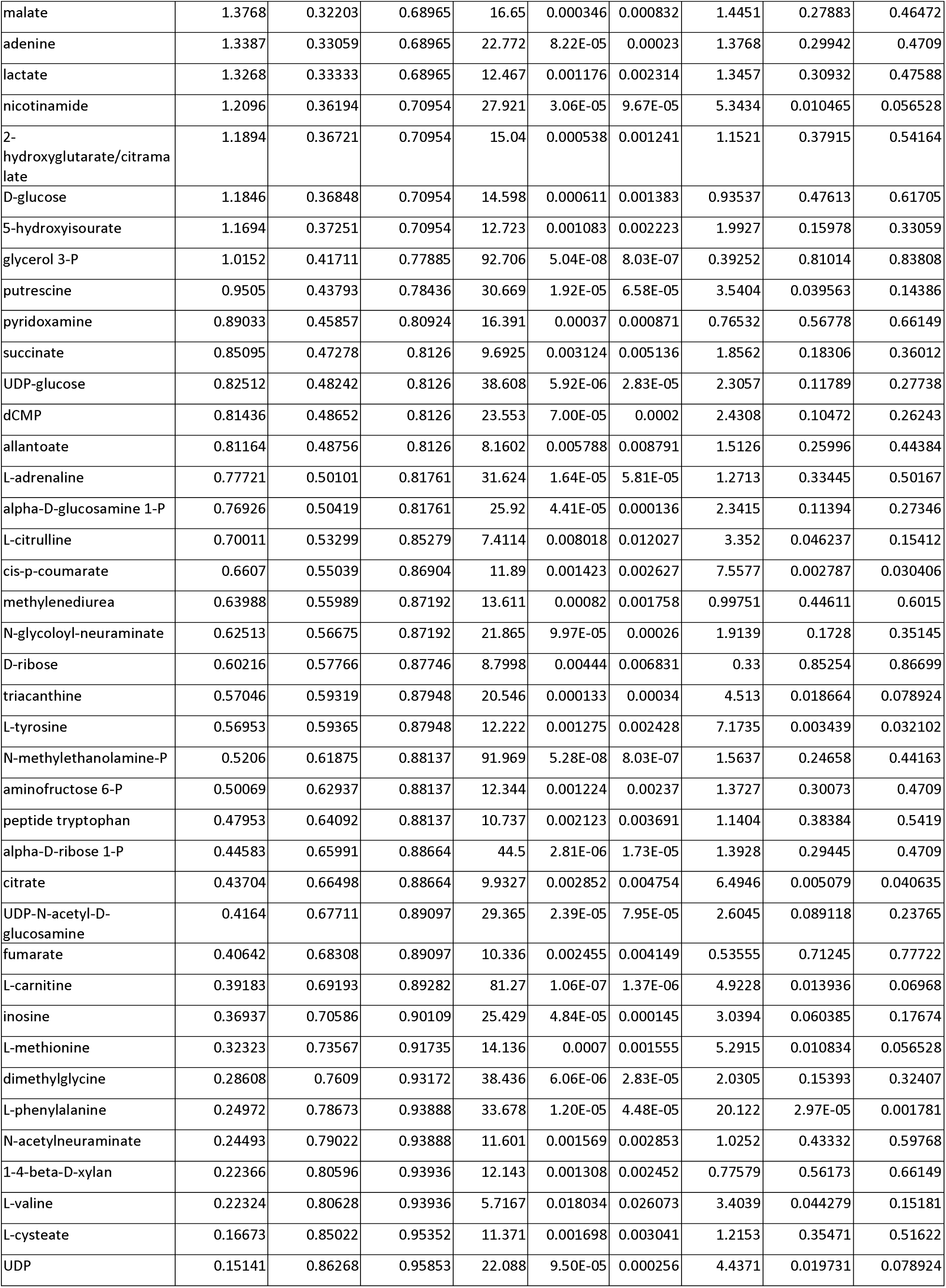
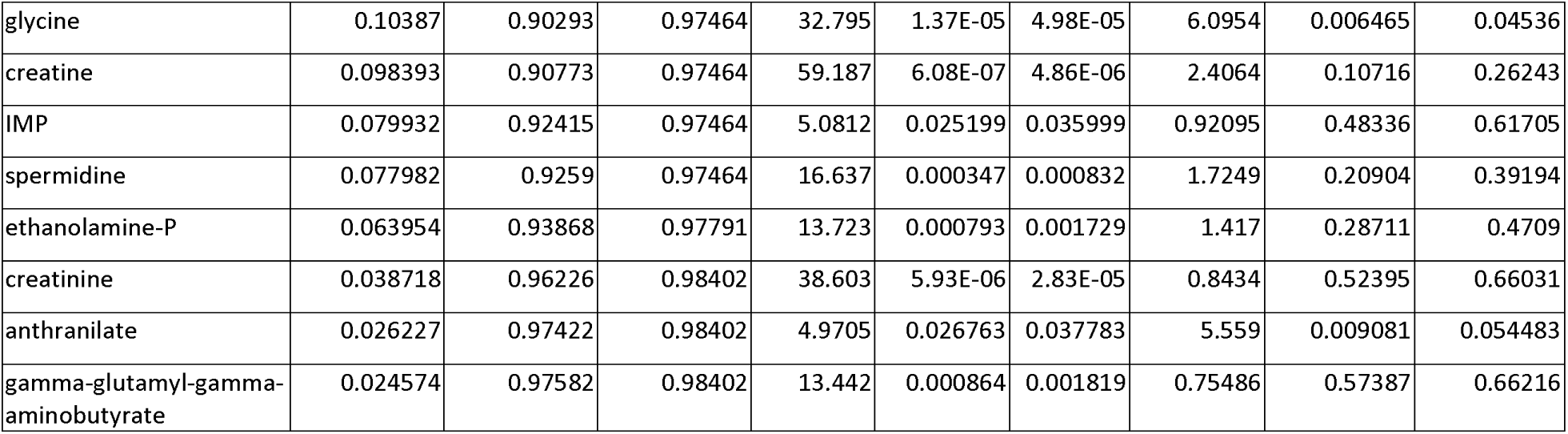
Two-way repeated measures analysis of variance (ANOVA).

**Supplemental Table 5.** Metabolic pathway analysis.

|  | Total | Expected | Hits | Raw p | -LOG(p) | Holm adj. | FDR | Impact |
| --- | --- | --- | --- | --- | --- | --- | --- | --- |
| Aminoacyl-tRNA biosynthesis | 75 | 0.6855 | 10 | 2.25E-10 | 22.215 | 1.80E-08 | 1.80E-08 | 0.16902 |
| Nitrogen metabolism | 39 | 0.35646 | 8 | 6.06E-10 | 21.225 | 4.79E-08 | 2.42E-08 | 0.0083 |
| Alanine, aspartate and glutamate metabolism | 24 | 0.21936 | 5 | 1.49E-06 | 13.416 | 0.000116 | 3.98E-05 | 0.69421 |
| Cyanoamino acid metabolism | 16 | 0.14624 | 4 | 8.88E-06 | 11.632 | 0.000684 | 0.000178 | 0 |
| Glycine, serine and threonine metabolism | 48 | 0.43872 | 5 | 5.21E-05 | 9.8624 | 0.003959 | 0.000834 | 0.42039 |
| Glutathione metabolism | 38 | 0.34732 | 4 | 0.000315 | 8.0617 | 0.023655 | 0.004205 | 0.24838 |
| Cysteine and methionine metabolism | 56 | 0.51184 | 4 | 0.001408 | 6.5656 | 0.10419 | 0.01609 | 0.03581 |
| Valine, leucine and isoleucine biosynthesis | 27 | 0.24678 | 3 | 0.001683 | 6.3874 | 0.12283 | 0.016827 | 0.03498 |
| D-Glutamine and D-glutamate metabolism | 11 | 0.10054 | 2 | 0.004174 | 5.4789 | 0.30052 | 0.036556 | 0.13904 |
| Arginine and proline metabolism | 77 | 0.70378 | 4 | 0.00457 | 5.3883 | 0.32444 | 0.036556 | 0.03582 |
| Phenylalanine metabolism | 45 | 0.4113 | 3 | 0.007337 | 4.9149 | 0.51357 | 0.053358 | 0.11906 |
| Glyoxylate and dicarboxylate metabolism | 50 | 0.457 | 3 | 0.009838 | 4.6215 | 0.67883 | 0.065587 | 0.01316 |
| Citrate cycle (TCA cycle) | 20 | 0.1828 | 2 | 0.013719 | 4.289 | 0.93287 | 0.078392 | 0.15351 |
| Taurine and hypotaurine metabolism | 20 | 0.1828 | 2 | 0.013719 | 4.289 | 0.93287 | 0.078392 | 0.35252 |
| Thiamine metabolism | 24 | 0.21936 | 2 | 0.019493 | 3.9377 | 1 | 0.10396 | 0 |
| Pantothenate and CoA biosynthesis | 27 | 0.24678 | 2 | 0.024384 | 3.7138 | 1 | 0.11475 | 0 |
| Phenylalanine, tyrosine and tryptophan biosynthesis | 27 | 0.24678 | 2 | 0.024384 | 3.7138 | 1 | 0.11475 | 0.008 |
| Methane metabolism | 34 | 0.31076 | 2 | 0.037502 | 3.2834 | 1 | 0.16668 | 0.01751 |
| Purine metabolism | 92 | 0.84088 | 3 | 0.049189 | 3.0121 | 1 | 0.20182 | 0.00791 |
| Butanoate metabolism | 40 | 0.3656 | 2 | 0.050455 | 2.9867 | 1 | 0.20182 | 0.08516 |
| Nicotinate and nicotinamide metabolism | 44 | 0.40216 | 2 | 0.059869 | 2.8156 | 1 | 0.21771 | 0 |
| Histidine metabolism | 44 | 0.40216 | 2 | 0.059869 | 2.8156 | 1 | 0.21771 | 0.00051 |
| Porphyrin and chlorophyll metabolism | 104 | 0.95056 | 3 | 0.066468 | 2.711 | 1 | 0.22434 | 0 |
| Primary bile acid biosynthesis | 47 | 0.42958 | 2 | 0.067302 | 2.6986 | 1 | 0.22434 | 0.01644 |
| Tyrosine metabolism | 76 | 0.69464 | 2 | 0.15164 | 1.8862 | 1 | 0.47028 | 0.04724 |
| Sulfur metabolism | 18 | 0.16452 | 1 | 0.15284 | 1.8784 | 1 | 0.47028 | 0 |
| Ether lipid metabolism | 23 | 0.21022 | 1 | 0.19117 | 1.6546 | 1 | 0.56642 | 0 |
| Sphingolipid metabolism | 25 | 0.2285 | 1 | 0.20603 | 1.5797 | 1 | 0.58865 | 0 |
| beta-Alanine metabolism | 28 | 0.25592 | 1 | 0.22783 | 1.4791 | 1 | 0.60094 | 0 |
| Glycolysis or Gluconeogenesis | 31 | 0.28334 | 1 | 0.24907 | 1.39 | 1 | 0.60094 | 0.0953 |
| Pentose phosphate pathway | 32 | 0.29248 | 1 | 0.25602 | 1.3625 | 1 | 0.60094 | 0 |
| Lysine biosynthesis | 32 | 0.29248 | 1 | 0.25602 | 1.3625 | 1 | 0.60094 | 0 |
| Vitamin B6 metabolism | 32 | 0.29248 | 1 | 0.25602 | 1.3625 | 1 | 0.60094 | 0.01914 |
| Pyruvate metabolism | 32 | 0.29248 | 1 | 0.25602 | 1.3625 | 1 | 0.60094 | 0.18254 |
| Terpenoid backbone biosynthesis | 33 | 0.30162 | 1 | 0.26291 | 1.3359 | 1 | 0.60094 | 0 |
| Ubiquinone and other terpenoid-quinone biosynthesis | 36 | 0.32904 | 1 | 0.28322 | 1.2615 | 1 | 0.62939 | 0 |
| Glycerophospholipid metabolism | 39 | 0.35646 | 1 | 0.303 | 1.194 | 1 | 0.65153 | 0.01641 |
| Valine, leucine and isoleucine degradation | 40 | 0.3656 | 1 | 0.30948 | 1.1729 | 1 | 0.65153 | 0.02232 |
| Folate biosynthesis | 42 | 0.38388 | 1 | 0.32225 | 1.1324 | 1 | 0.66104 | 0.03372 |
| Ascorbate and aldarate metabolism | 45 | 0.4113 | 1 | 0.341 | 1.0759 | 1 | 0.682 | 0.01617 |
| Lysine degradation | 47 | 0.42958 | 1 | 0.35322 | 1.0407 | 1 | 0.68921 | 0 |
| Pentose and glucuronate interconversions | 53 | 0.48442 | 1 | 0.3886 | 0.9452 | 1 | 0.74019 | 0 |
| Pyrimidine metabolism | 60 | 0.5484 | 1 | 0.42754 | 0.8497 | 1 | 0.79543 | 0 |

**Supplemental Table 6.**
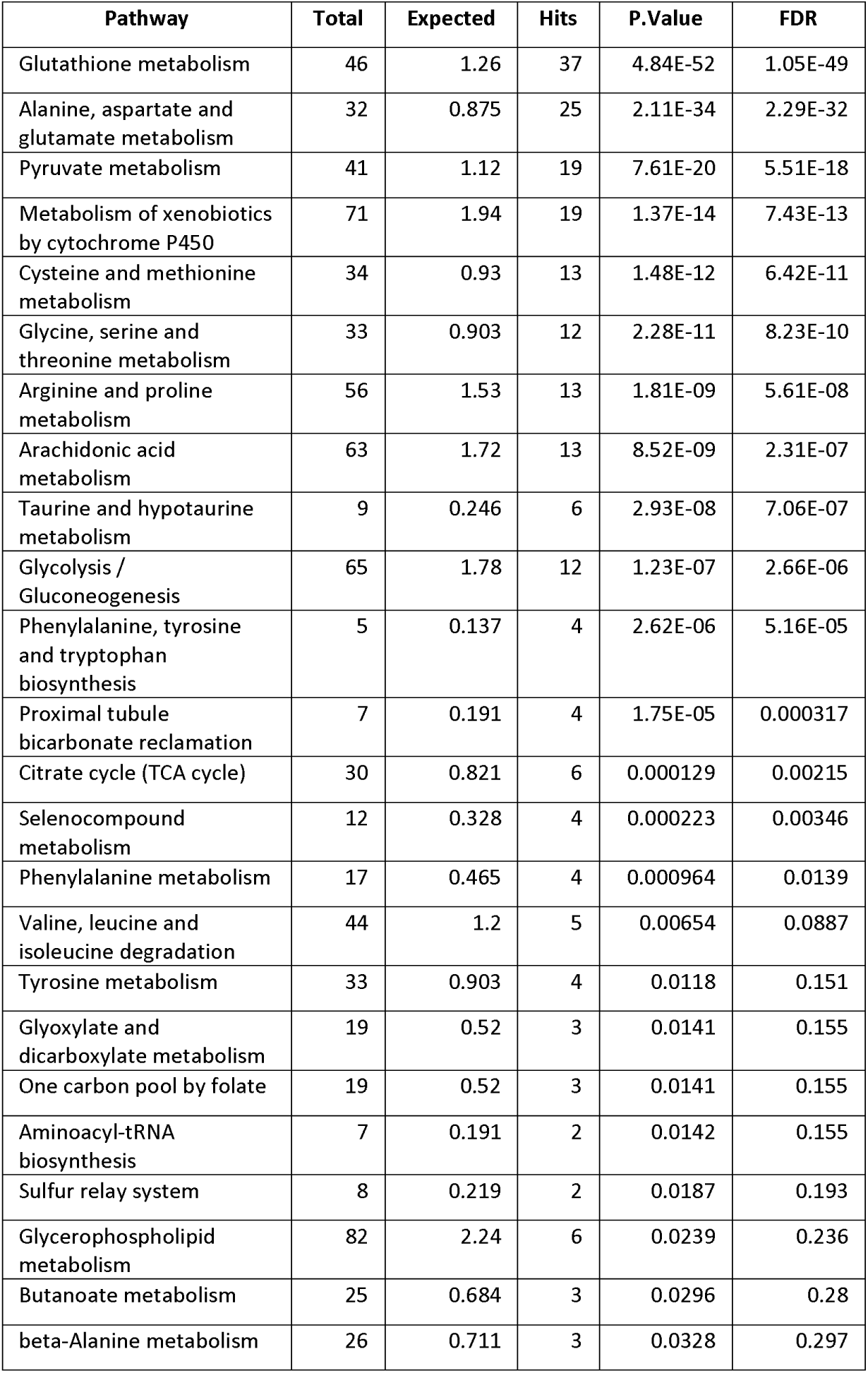
Metabolite-protein interaction nodal analysis.

| Pathway | Total | Expected | Hits | P.Value | FDR |
| --- | --- | --- | --- | --- | --- |
| Glutathione metabolism | 46 | 1.26 | 37 | 4.84E-52 | 1.05E-49 |
| Alanine, aspartate and glutamate metabolism | 32 | 0.875 | 25 | 2.11E-34 | 2.29E-32 |
| Pyruvate metabolism | 41 | 1.12 | 19 | 7.61E-20 | 5.51E-18 |
| Metabolism of xenobiotics by cytochrome P450 | 71 | 1.94 | 19 | 1.37E-14 | 7.43E-13 |
| Cysteine and methionine metabolism | 34 | 0.93 | 13 | 1.48E-12 | 6.42E-11 |
| Glycine, serine and threonine metabolism | 33 | 0.903 | 12 | 2.28E-11 | 8.23E-10 |
| Arginine and proline metabolism | 56 | 1.53 | 13 | 1.81E-09 | 5.61E-08 |
| Arachidonic acid metabolism | 63 | 1.72 | 13 | 8.52E-09 | 2.31E-07 |
| Taurine and hypotaurine metabolism | 9 | 0.246 | 6 | 2.93E-08 | 7.06E-07 |
| Glycolysis / Gluconeogenesis | 65 | 1.78 | 12 | 1.23E-07 | 2.66E-06 |
| Phenylalanine, tyrosine and tryptophan biosynthesis | 5 | 0.137 | 4 | 2.62E-06 | 5.16E-05 |
| Proximal tubule bicarbonate reclamation | 7 | 0.191 | 4 | 1.75E-05 | 0.000317 |
| Citrate cycle (TCA cycle) | 30 | 0.821 | 6 | 0.000129 | 0.00215 |
| Selenocompound metabolism | 12 | 0.328 | 4 | 0.000223 | 0.00346 |
| Phenylalanine metabolism | 17 | 0.465 | 4 | 0.000964 | 0.0139 |
| Valine, leucine and isoleucine degradation | 44 | 1.2 | 5 | 0.00654 | 0.0887 |
| Tyrosine metabolism | 33 | 0.903 | 4 | 0.0118 | 0.151 |
| Glyoxylate and dicarboxylate metabolism | 19 | 0.52 | 3 | 0.0141 | 0.155 |
| One carbon pool by folate | 19 | 0.52 | 3 | 0.0141 | 0.155 |
| Aminoacyl-tRNA biosynthesis | 7 | 0.191 | 2 | 0.0142 | 0.155 |
| Sulfur relay system | 8 | 0.219 | 2 | 0.0187 | 0.193 |
| Glycerophospholipid metabolism | 82 | 2.24 | 6 | 0.0239 | 0.236 |
| Butanoate metabolism | 25 | 0.684 | 3 | 0.0296 | 0.28 |
| beta-Alanine metabolism | 26 | 0.711 | 3 | 0.0328 | 0.297 |

**Supplemental Table 7.**
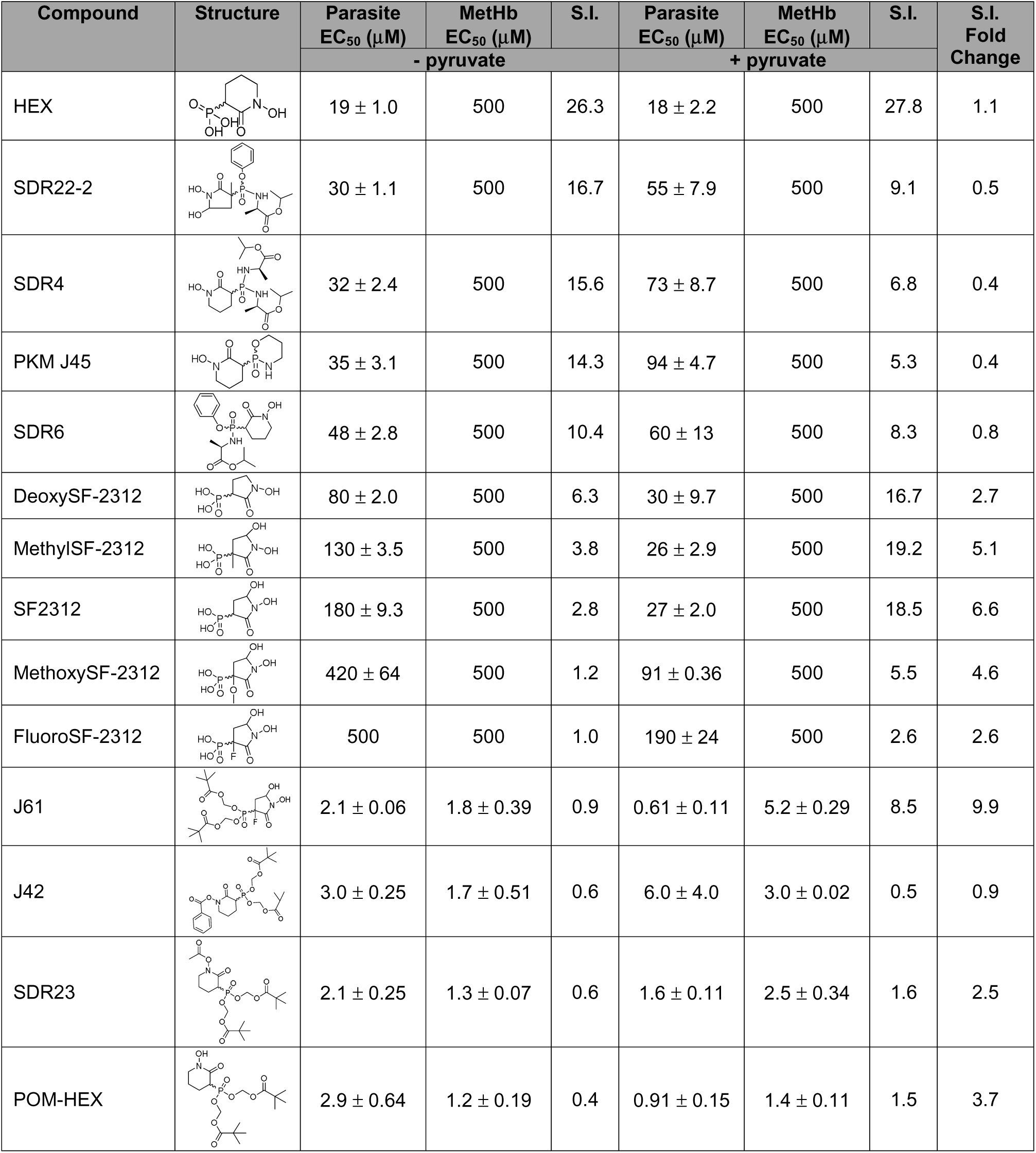
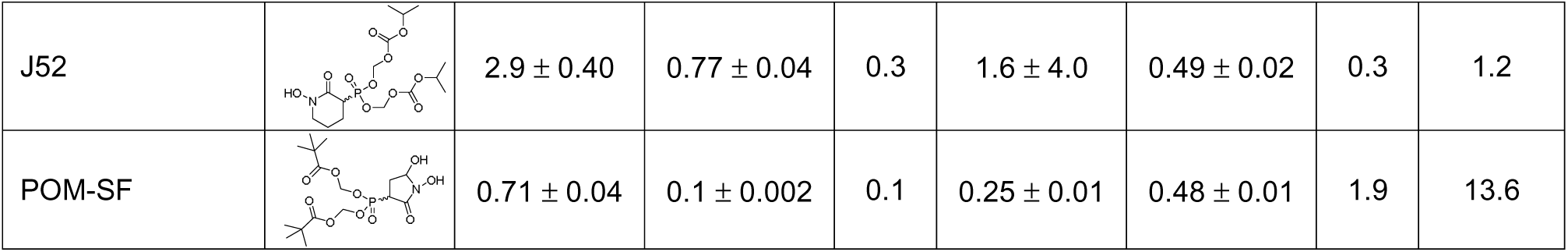
Table of parasite EC_50_s and methemoglobin EC_50_s for all tested compounds and their structures.

| Compound | Structure | Parasite<br>EC<sub>50</sub> (μM) | MetHb<br>EC<sub>50</sub> (μM) | S.I. | Parasite<br>EC<sub>50</sub> (μM) | MetHb<br>EC<sub>50</sub> (μM) | S.I. | S.I.<br>Fold<br>Change |
| --- | --- | --- | --- | --- | --- | --- | --- | --- |
|  |  | - pyruvate |  |  | + pyruvate |  |  |  |
| HEX |  | 19 ± 1.0 | 500 | 26.3 | 18 ± 2.2 | 500 | 27.8 | 1.1 |
| SDR22-2 |  | 30 ± 1.1 | 500 | 16.7 | 55 ± 7.9 | 500 | 9.1 | 0.5 |
| SDR4 |  | 32 ± 2.4 | 500 | 15.6 | 73 ± 8.7 | 500 | 6.8 | 0.4 |
| PKM J45 |  | 35 ± 3.1 | 500 | 14.3 | 94 ± 4.7 | 500 | 5.3 | 0.4 |
| SDR6 |  | 48 ± 2.8 | 500 | 10.4 | 60 ± 13 | 500 | 8.3 | 0.8 |
| DeoxySF-2312 |  | 80 ± 2.0 | 500 | 6.3 | 30 ± 9.7 | 500 | 16.7 | 2.7 |
| MethylSF-2312 |  | 130 ± 3.5 | 500 | 3.8 | 26 ± 2.9 | 500 | 19.2 | 5.1 |
| SF2312 |  | 180 ± 9.3 | 500 | 2.8 | 27 ± 2.0 | 500 | 18.5 | 6.6 |
| MethoxySF-2312 |  | 420 ± 64 | 500 | 1.2 | 91 ± 0.36 | 500 | 5.5 | 4.6 |
| FluoroSF-2312 |  | 500 | 500 | 1.0 | 190 ± 24 | 500 | 2.6 | 2.6 |
| J61 |  | 2.1 ± 0.06 | 1.8 ± 0.39 | 0.9 | 0.61 ± 0.11 | 5.2 ± 0.29 | 8.5 | 9.9 |
| J42 |  | 3.0 ± 0.25 | 1.7 ± 0.51 | 0.6 | 6.0 ± 4.0 | 3.0 ± 0.02 | 0.5 | 0.9 |
| SDR23 |  | 2.1 ± 0.25 | 1.3 ± 0.07 | 0.6 | 1.6 ± 0.11 | 2.5 ± 0.34 | 1.6 | 2.5 |
| POM-HEX |  | 2.9 ± 0.64 | 1.2 ± 0.19 | 0.4 | 0.91 ± 0.15 | 1.4 ± 0.11 | 1.5 | 3.7 |
| J52 | | $2.9 \pm 0.40$ | $0.77 \pm 0.04$ | 0.3 | $1.6 \pm 4.0$ | $0.49 \pm 0.02$ | 0.3 | 1.2 |
| POM-SF | | $0.71 \pm 0.04$ | $0.1 \pm 0.002$ | 0.1 | $0.25 \pm 0.01$ | $0.48 \pm 0.01$ | 1.9 | 13.6 |

**Supplemental Table 8.** Enolase inhibitors are active against multidrug-resistant *P. falciparum*. Half-maximal inhibitor concentrations (EC_50_s) were determined for drug resistant malaria parasites and compared to the sensitive 3D7 strain. The respective EC_50_s are calculated from each of the independent biological replicates using a non-linear regression of the log of the inhibitor concentration with data from each technical replicate normalized to maximal and minimal growth using the software package GraphPad Prism. Displayed are the means ± s.e.m calculated from three biological replicates.

| Cell line | POM-SF EC<sub>50</sub> (nM) | POM-HEX EC<sub>50</sub> (nM) |
| --- | --- | --- |
| *P. falciparum* 3D7<br>(pan-sensitive) | 185 ± 18 | 910 ± 148 |
| *P. falciparum* K1<br>(chloroquine and sulfadoxine-pyrimethamine resistant) | 361 ± 34 | 1112 ± 530 |
| *P. falciparum* D10<br>(mefloquine resistant) | 437 ± 83 | 1432 ± 540 |
| *P. falciparum* IPC-5202<br>(chloroquine and artemisinin<br>resistant) | 321 ± 11 | 966 ± 442 |

